# Tumor stiffening reversion through collagen crosslinking inhibition improves T cell migration and anti-PD-1 treatment

**DOI:** 10.1101/2020.05.19.104430

**Authors:** Alba Nicolas-Boluda, Javier Vaquero, Sarah Barrin, Chahrazade Kantari-Mimoun, Matteo Ponzo, Gilles Renault, Piotr Deptuła, Katarzyna Pogoda, Robert Bucki, Ilaria Cascone, Jose Courty, Laura Fouassier, Florence Gazeau, Emmanuel Donnadieu

## Abstract

Only a fraction of cancer patients benefits from immune checkpoint inhibitors. This may be partly due to the dense extracellular matrix (ECM) that forms a barrier for T cells. Comparing 5 preclinical mouse tumor models with heterogeneous tumor microenvironments, we aimed to relate the rate of tumor stiffening with the remodeling of ECM architecture and to determine how these features affect intratumoral T cell migration. An ECM-targeted strategy, based on the inhibition of lysyl oxidase (LOX) was used. In vivo stiffness measurements were found to be strongly correlated with tumor growth and ECM crosslinking but negatively correlated with T cell migration. Interfering with collagen stabilization reduces ECM content and tumor stiffness leading to improved T cell migration and increased efficacy of anti-PD-1 blockade. This study highlights the rationale of mechanical characterizations in solid tumors to understand resistance to immunotherapy and of combining treatment strategies targeting the ECM with anti-PD-1 therapy.

## INTRODUCTION

In the last decade, a significant progress has been made in the development of T cell-based immunotherapies (*1*). The two main T-cell based immunotherapies are adoptive T cell therapy and immune checkpoint inhibitors. Monoclonal antibodies blocking the immune checkpoints cytotoxic T lymphocyte associated antigen 4 (CTLA-4) and programmed death 1 receptor (PD-1) have quickly gone from proof of concepts to FDA approved first and second line treatments for a significant number of tumors even in late stages (*2*). However, an elevated percentage of patients with solid tumors fail to respond to these therapies. The mechanisms underlying the poor response to immune checkpoint inhibitors are still uncertain, nevertheless recent results suggest that T cell function and distribution in the tumor are affected by numerous immunosuppressive mechanisms (*3*). It is well established that in progressing tumors T cells exhibit a particular phenotype unable to normally respond to tumor antigens. In addition, in a large proportion of tumors, T lymphocytes are excluded from the tumor cell regions in a so called “excluded-immune profile” (*4-6*). Ineffective T cell migration and penetration into the tumor mass might represent an important obstacle to T cell-based immunotherapies. As a support for this notion, various clinical studies have shown that tumors enriched in T cells are more susceptible to be controlled by PD-1 blockade. In contrast, tumors with so-called immune excluded profiles, in which T cell are present within tumors but not in contact with malignant cells, are refractory to PD-1 blockade (*4, 7*). Particularly, the fibrotic state of desmoplastic tumors can cause immunosuppression through multiple mechanisms (*8*). The hypothesis of a physical resistance to T cell infiltration and migration related to the heterogeneity and aberrant organization of the extracellular matrix with respect to the tumor nests has emerged recently (*9, 10*). By using dynamic imaging microscopy, we highlighted the detrimental impact of collagen fibrils architecture on the migratory behavior of T cells in fresh human tumor explants. Both a guiding strategy combined to a physical hindrance process have been shown to restrain T cells from contacting tumor cells, thus leading to the T cell excluded profile (*11, 12*). Hence, a dense fibrotic stroma could raise physical obstacles to immune cell infiltration similar to the previously established stromal resistance to chemotherapeutics, antibodies, nanoparticles or virus tumor penetration (*13, 14*). In addition, cellular components of tumor-associated fibrosis, particularly the cancer-associated fibroblasts (CAF), can have both direct and indirect effects on T cell infiltration and function (*8*). Accordingly, one important challenge in the field is to develop strategies targeting tumor fibrosis in order to reverse immune exclusion and to improve T cell-based immunotherapy. Recent studies have been undertaken with this objective. T cells engineered to express a chimeric antigen receptor together with heparanase, an ECM degrading enzyme, show enhanced infiltration into xenografted tumors as well as anti-tumor efficacy (*15*). Recently, a major role for the TGFβ signaling pathway in promoting T cell exclusion from tumor cells has been demonstrated. In breast mouse tumor models, neutralizing antibodies against TGFβ were shown to reduce collagen I production, overcoming the T cell excluded profile and increasing the efficacy of anti-PD-L1 antibodies (*7, 16, 17*). In cholangiocarcinoma, an immune mesenchymal subtype has been identified, which is associated with TGFβ signature and poor tumor infiltrating cells (*18*). Other axes including the CXCR4/CXCL12 in breast metastasis and the focal adhesion kinase in pancreatic ductal adenocarcinoma (PDAC) have also been associated with both desmoplasia and absence of cytotoxic T lymphocytes in tumors from mice (*19*). Consequently, the inhibition of these axes in preclinical mouse cancer models was shown to reduce fibrosis while significantly increasing T cell infiltration and improving response to checkpoint inhibitors (*7, 16, 20, 21*). Clinical trials testing such combination are currently ongoing in advanced pancreatic cancer, mesothelioma, urethelial carcinoma and other malignancies (NCT02546531, NCT02758587, NCT02734160, NCT04064190, NCT02947165).

However due to patient and tumor heterogeneity, there is no clear indication on how the T cell distribution in tumors is related to the fibrosis level and to the different architectures of ECM. Thus, there is an urgent need to assist in matching combination approaches to patient populations who could benefit from stromal modulation strategies to improve their response to immunotherapy. Companion matrix-derived biomarkers and imaging approaches should provide insights into the contribution of the ECM remodeling in shaping the immune milieu of the tumor. Particularly, a critical determinant of fibrotic tumor progression - the tumor mechanics - has been poorly investigated through the prism of immune impact. An important feature of fibrotic tumors is their considerable higher stiffness compared to their neighboring healthy tissues which is highly correlated with cancer progression and metastasis, particularly in breast, colorectal, liver and pancreatic tumors (*22, 23*). The use of non-invasive imaging techniques such as shear wave elastography (SWE) and magnetic resonance elastography (MRE), designed to monitor stiffness of any given tissue, allow an accurate diagnostic and characterization of malignant lesions *in vivo* (*24*). The extensive remodeling of the stromal components increasing tumor stiffness can mechanically activate intracellular signaling pathways that promote tumor progression and at the same time can dampen T cell functions including migration and infiltration into tumor islets (*25-28*). However, there is a lack of studies correlating the mechanical properties of tumors to their heterogeneous ECM architecture and T cell infiltration capacity. Here we aim at filling this gap through a comprehensive investigation of stiffness evolution in several preclinical mouse models of pancreatic, breast and bile duct carcinomas, presenting different ECM organization, coupled to dynamic imaging of fresh tumor slices to monitor T cell motility. In concert with these imaging biomarkers of both mechanical properties, ECM architecture and T cell migration, we explored the consequences of altering the ECM by inhibition of the lysyl oxidase (LOX), a copper-dependent enzyme responsible for the crosslinking of collagen molecules into fibers that has been seen to be overexpressed in many metastatic tumors and responsible for malignant progression (*29, 30, 31*). We highlight that LOX inhibition has different mechanical modulating effects depending on the ECM architecture, with significant improvement in T cell mobility. Despite minor effects in primary tumor growth upon LOX inhibition or PD-1 blockade treatment alone, their combination increases effector CD8 T cell accumulation in tumors and significantly delays tumor progression in a pancreatic cancer model.

## RESULTS

### Relationship between tumor structure and tumor mechanical properties in different preclinical carcinoma mouse models

One key aspect when testing immunotherapeutic agents is the use of relevant preclinical models that closely mimic the properties of human solid tumors. Human carcinomas derive from epithelial cells and therefore harbor a typical though heterogeneous structure with tumor cells forming compact islets or nests surrounded by the stroma, enriched in ECM proteins, fibroblasts, blood vessels, and immune cells. To unravel the relationship between tumor growth, ECM remodeling, stiffening and immune infiltration, we characterized the tumor structure and the mechanical properties of five different preclinical models, recapitulating the structure heterogeneity of different carcinomas (**Table S1**): subcutaneous model of cholangiocarcinoma (EGI-1), subcutaneous (MET-1) and transgenic model (MMTV-PyMT) of mouse breast carcinoma, orthotopic (mPDAC) and subcutaneous (KPC) models of mouse pancreatic ductal adenocarcinoma. A multiscale evaluation of the mechanical properties of the tumors was performed. At the macroscale, we measured tumor stiffness during tumor growth using SWE, a noninvasive imaging technique that allows the quantification and mapping of tumor stiffness (**Figure 1A, Table 1**). The presence of very stiff regions, defined as areas with an elastic modulus > 40 kPa, in the tumor was quantified together with the average stiffness of the tumor (**Figure 1B, Table 1**). At the micron-scale, we evaluated tumor organization and fibrosis using hematoxylin-eosin-safran (HES) (**Figure 1C**) and Sirius Red staining. Sirius Red is a highly specific stain for collagen fibers that combined with polarizing microscopy allows differentiating thin collagen fibrils from thick and densely packed collagen fibers (*32*). Under polarized light, thin fibers show a greenish-yellow birefringence whilst thicker and densely packed fibers give an orange red birefringence. By separating these two colors, it is possible to quantify the amount of thick and densely packed fibers present in the tumor (**Table 1**). The fibrillar collagen network was determined using second harmonic generation imaging (SHG), that allows to analyze the architecture and density of fibrillar collagen without having to use detection antibodies (**Figure 1D, Table 1**).

**Table 1.** Tumor stiffness and tumor architecture parameters for EGI-1, KPC, MMTV-PyMT, mPDAC and MET-1 models. Mean stiffness of tumors with volume > 600 mm^3^ measured with SWE. Percentage of stiff regions of tumors with volume > 600 mm^3^. Tumor architecture was characterized by the percentage of the tumor covered by the stromal compartment, estimated from HES images; collagen fiber width and length, calculated from SHG images; percentage of red-orange birefringent fibers combining red Sirius staining and polarized microscopy, orange-red fibers correspond to thick and packed regions.

| Tumor model | Tumor Stiffness at > 600 m<sup>3</sup> |  |  | Tumor Architecture |  |  |
| --- | --- | --- | --- | --- | --- | --- |
|  | Mean stiffness (kPa) | % Tumor area > 40 kPa | % Area stroma | Collagen fiber width (μm) | Collagen fiber length (μm) | % Sirius red staining |
| EGI-1 | 29.9 ± 10.0 | 19.2 ± 10.0 | 20.8 ± 5.6 | 7.44 ± 2.2 | 85.9 ± 42.0 | 6.0 ± 5.0 |
| KPC | 26.4 ± 8.0 | 21.2 ± 16.0 | 19.3 ± 6.9 | 4.48 ± 0.9 | 67.1 ± 35.0 | 4.1 ± 1.4 |
| MMTV-PyMT | 29.7 ± 7.0 | 15.4 ± 12.0 | 13.1 ± 6.9 | 3.66 ± 1.1 | 83.8 ± 41.0 | 4.8 ± 3.5 |
| mPDAC | 13.6 ± 6.0 | 0.4 ± 0.3 | 43.4 ± 5.5 | 3.49 ± 0.8 | 81.5 ± 38.0 | 2.7 ± 2.1 |
| MET-1 | 10.3 ± 4.0 | 0.2 ± 0.1 | 6.13 ± 3.1 | 3.21 ± 0.9 | 73.8 ± 31.0 | - |

**Figure 1.**
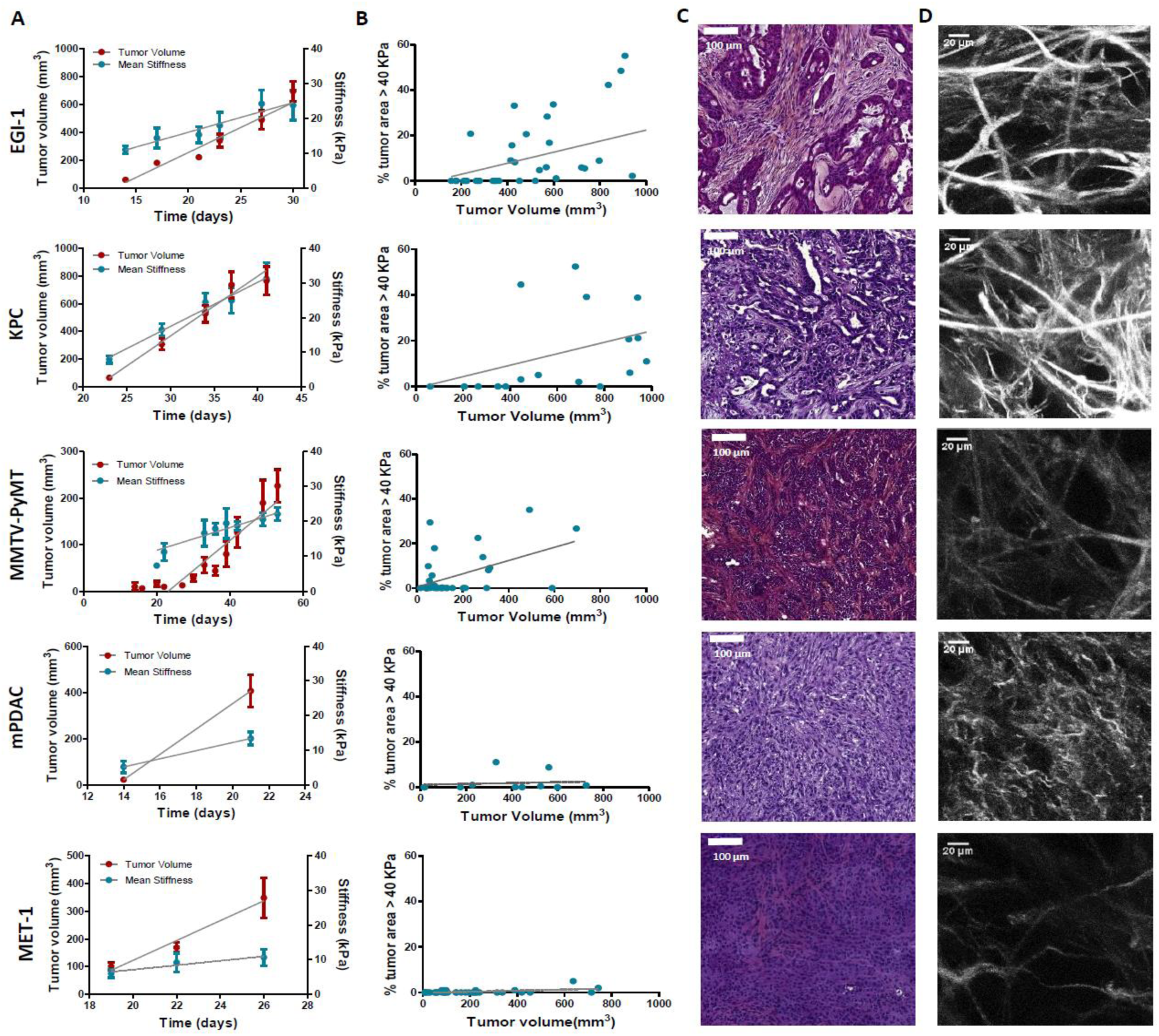
Macroscopic and microscopic characterization of EGI-1, KPC, MMTV-PyMT, mPDAC, MET-1 tumor models. (**A**) Tumor volume (right Y axis) and tumor mean stiffness (left Y axis) evolution in time. Tumor volume was measured using a caliper or ultrasound, whilst tumor mean stiffness was measured using SWE. (**B**) The percentage of stiff regions in relation to tumor volume. The stiff regions were defined as regions with an elastic modulus > 40 kPa. (**C**) Comparison of the histological diversity within the tumor models. Representative images of each tumor model (Scale bar = 100 µm). (**D**) SHG images of collagen network in each of the models at the endpoint of the experiment. Scale bar = 20 µm.

In the EGI-1 cholangiocarcinoma model, tumor stiffening and tumor growth have a strong positive correlation (**Figure 1A**). The stiffness distribution is highly heterogeneous, presenting 20 % of stiff regions (> 40 kPa) in average that goes up to 50-60 % in tumors with higher volume. In terms of architecture, the tumor and its extensive stroma compartment, occupying around 20 % of the tumor, are well separated. This is a typical trait of desmoplastic tumors and the model accurately reproduces the architecture of human cholangiocarcinoma. Its collagen network is characterized by long (85.9 ± 42.0 µm) and thick (7.4 ± 2.2 µm) collagen fibers that are densely packed (6 % of the tumor) (**Table 1, Figure 1D**).

The mouse pancreatic ductal adenocarcinoma KPC model (*33*) exhibits similar features to that of EGI-1. A high positive correlation between tumor stiffness and volume, presenting over 20 % of stiff regions at high tumor volumes, was observed (**Figure 1A**). KPC tumors present a typical segregation of ECM and tumor nests, with a high proportion of stroma (∼20 %). However, the extension of the stromal areas is lower than that of EGI-1 with higher intercalation with tumor islets. Its collagen network is characterized by shorter (67.1 ± 35 µm) and thinner (4.4 ± 0.9 µm) collagen fibers than that of EGI-1 (**Table1, Figure 1D**).

The spontaneous orthotopic murine breast cancer model MMTV PyMT, although slower in its growth as compared to subcutaneous tumors also stiffens during tumor progression (**Figure 1A**). However, there is a lower density of stiff regions (<16%). Of note, the fact that it is an orthotopic tumor in a genetic model that develops about 10 tumors limits the maximal tumor volume reached for this analysis. Hence, we cannot compare this model with the other models at high tumor volumes for ethical reasons. This spontaneous tumor model also presents a tumor islets-stroma structure, but with a lower amount of stroma (∼13 %) as compared to EGI-1 and KPC models. The stroma is more dispersed and intercalated with the tumor compartment. Collagen fibers are characterized for being thin (3.6 ± 1.1 µm) and long (83 ± 41 µm), forming densely packed regions taking up to 4.8 % of the tumor.

The orthotopic murine pancreatic ductal adenocarcinoma (mPDAC) model has a very different profile compared to the other models. The stroma takes up ∼ 40 % of the tumor, however it is well nested into the tumor with no spatial segregation of stromal and tumor compartment. mPDAC cells have already undergone the epithelial mesenchymal transition (EMT), which can explain this architecture (*34*). Cells that have undergone EMT gain the capacity of secreting ECM proteins that modulate the structure of the stroma. The mPDAC collagen network is made up of thin (3.5 ± 0.8 µm) and dispersed collagen fibers, which accounts for the lower presence of densely packed collagen regions (2.7 % of the tumor). The mean tumor stiffness is lower than that of the other models, partly explained by the limits of the maximal tumor volume reached in this orthotopic model (for ethical reasons) and partly, to the collagen architecture. Tumor stiffness also increases with tumor volume (**Figure 1B**) in line with previous studies performed in this model (*34*). Unlike the above mentioned models which exhibit a high correlation between stiffness and tumor growth, the mouse breast carcinoma MET-1 tumor model is characterized by low tumor stiffness (**Figure 1A**), a limited stroma (∼ 6 %), the lack of tumor-islet/stroma organization and the presence of thin (3.2 ± 0.9 µm) and dispersed collagen fibers (**Table 1, Figure 1D**).

This thorough analysis enabled us to confirm the correlation of high tumor stiffness measured non-invasively with collagen accumulation associated to a segregated architecture of thick and densely packed collagen fibers (Sirius red positive) surrounding tumor nests. In contrast, tumors with entangled and thin mesh of collagen present lower rigidity despite overall high collagen content. Particularly the appearance of stiff regions > 40kPa is seen as a physical biomarker of intratumor heterogeneity and ECM segregation. This analysis maps out potentially relevant preclinical tumor models which might reproduce the fibrotic evolution of human breast, pancreatic and bile duct tumors and their architecture heterogeneity.

### LOX modulates tumor stiffness and the ECM organization

The panel of tumor stroma structures reported above allows us to investigate the direct effects of ECM modulating agents in situations mimicking the heterogeneity observed in human carcinoma. Thus we sought to determine whether beta-aminopropionitrile (BAPN), an inhibitor for LOX enzymatic activity, could modulate tumors’ mechanical properties in concert with the stroma architecture (*31*). For these experiments, BAPN was administered in the drinking water of mice upon tumor cell implantation and until their sacrifice for most models, except for MMTV-PyMT model that was treated approximately at the time that tumors start to spontaneously develop. We first examined the effect of LOX inhibition on tumor stiffness (**Figures 2A and 2C**) and on the presence of stiff regions (**Figure 2B**). Results show that all models, except for MET-1, undergo reduction in mean stiffness when LOX is inhibited. EGI-1 and KPC models both show the most striking differences. The change is mainly perceived at late stages of tumor development, since in these models, tumor stiffness is positively correlated with tumor growth. In the MMTV-PyMT model, however, significant differences were noted throughout the development of the tumor. For the mPDAC model, tumor stiffness was only evaluated at the end of the BAPN treatment. A significant decrease of mean tumor stiffness is seen in BAPN treated mPDAC tumors.

**Figure 2.**
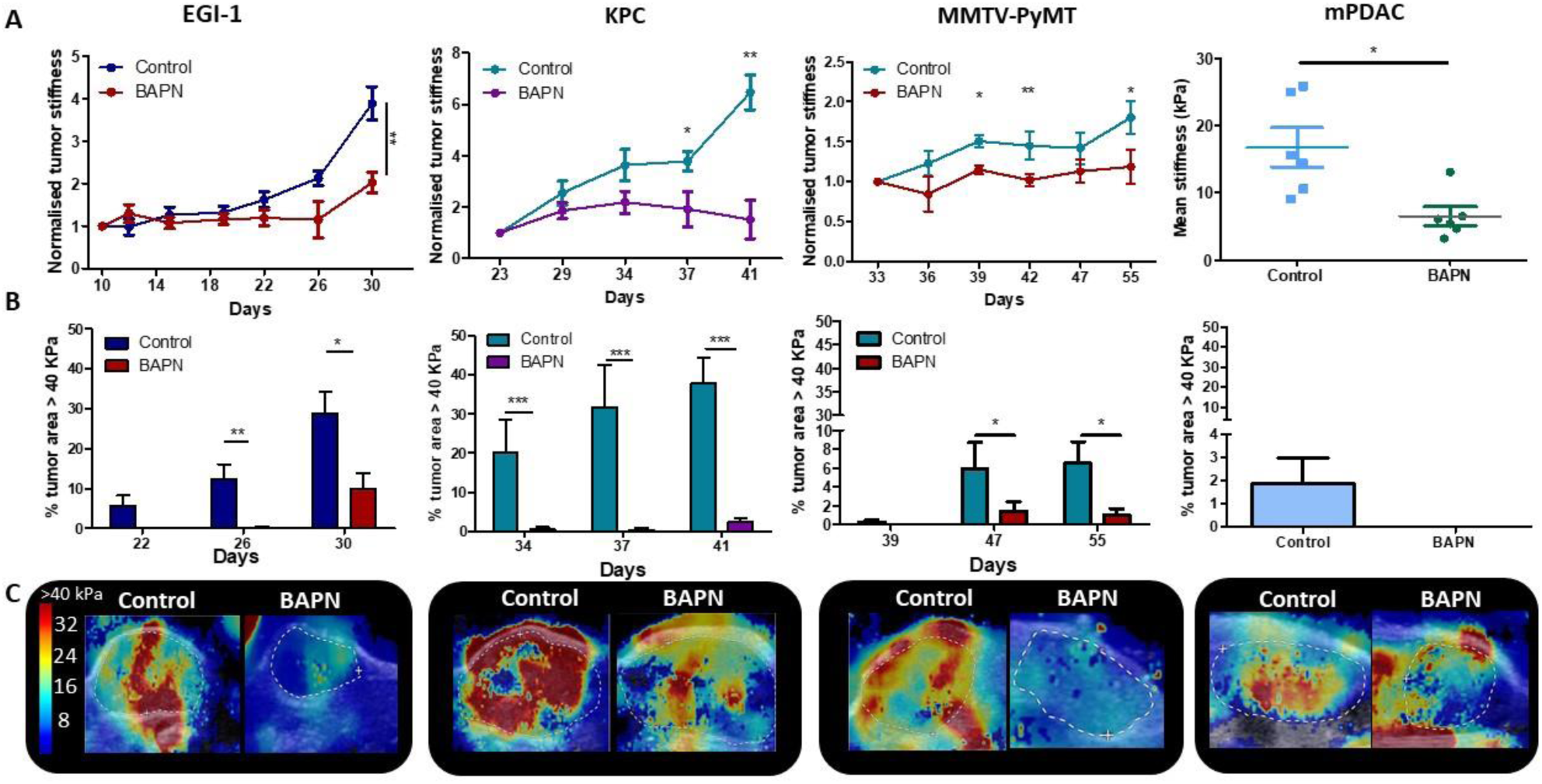
Effect of LOX inhibition on tumor stiffness and tumor stroma organization in EGI-1, KPC, MMTV-PyMT and mPDAC tumor models. (**A**) Tumor stiffness measured by SWE in control and BAPN treated (LOX inhibitor) tumors in relation to tumor volume for EGI-1, MMTV-PyMT and KPC tumor models. For mPDAC tumor stiffness was only measured at the endpoint of the experiment. (\**P-* value<0.05, \*\**P-*value<0.01, \*\*\**P-*value<0.001 Students’ t test). (**B**) Percentage of tumor area with stiffness > 40kPa at 3 different time points in the late-stages of tumor development. (**C**) Illustrative examples of SWE images of each on the tumors at the last time point of the experiment (EGI-1 – day 30, KPC – day 41, MMTV-PyMT – day 55, mPDAC – day 21).

Notably, BAPN treatment did not affect tumor growth in most models (**Figures S1A-S5A**), except for mPDAC (**Figure S4A**). To verify that the variation in tumor stiffness was not due to difference in tumor volume, the mean tumor stiffness of control and treated mice were compared at different tumor volumes (**Figures S2C-S6C**). In both KPC and EGI-1 tumors a clear difference in mean tumor stiffness can be seen in tumors with a volume > 400 mm^3^ (**Figures S1C and S2C**).

We also explored whether the presence and proportion of stiff regions was reduced when LOX was inhibited (**Figure 2B**). The percentage of control tumor area with a mean stiffness >40kPa increased with time (and tumor volume), indicating that there is not only an increase of overall mean stiffness, but also an increase of the heterogeneity of stiff regions within the non-treated tumor. However, this percentage was significantly reduced in BAPN-treated tumors with marked differences observed in the KPC model and to a lesser extent in EGI-1 and MMTV-PyMT models. In mPDAC, BAPN-treated tumors did not display stiff regions (**Figure S4 C-E**). The only model that does not respond to LOX inhibition by stiffness reduction is MET-1 (**Figure S5 B-D**), in line with our previous data showing an absence of tumor stiffening during tumor growth. Overall, our results clearly illustrate: 1) the heterogeneity of tumor response to an ECM-targeting agent, 2) the potential of non-invasive SWE elastography to measure a macroscopic physical marker - stiffness-that predicts this response.

Given the effects of LOX inhibition at a macro scale, we decided to delve into the changes induced at the level of the collagen fiber network through an in-depth quantitative evaluation of collagen fiber width (**Figure 3A**), orientation (**Figures 3B and 3C**), curvature (**Figure 3D**) and the presence of regions with thick and densely packed fibers (**Figure 3E**).

**Figure 3.**
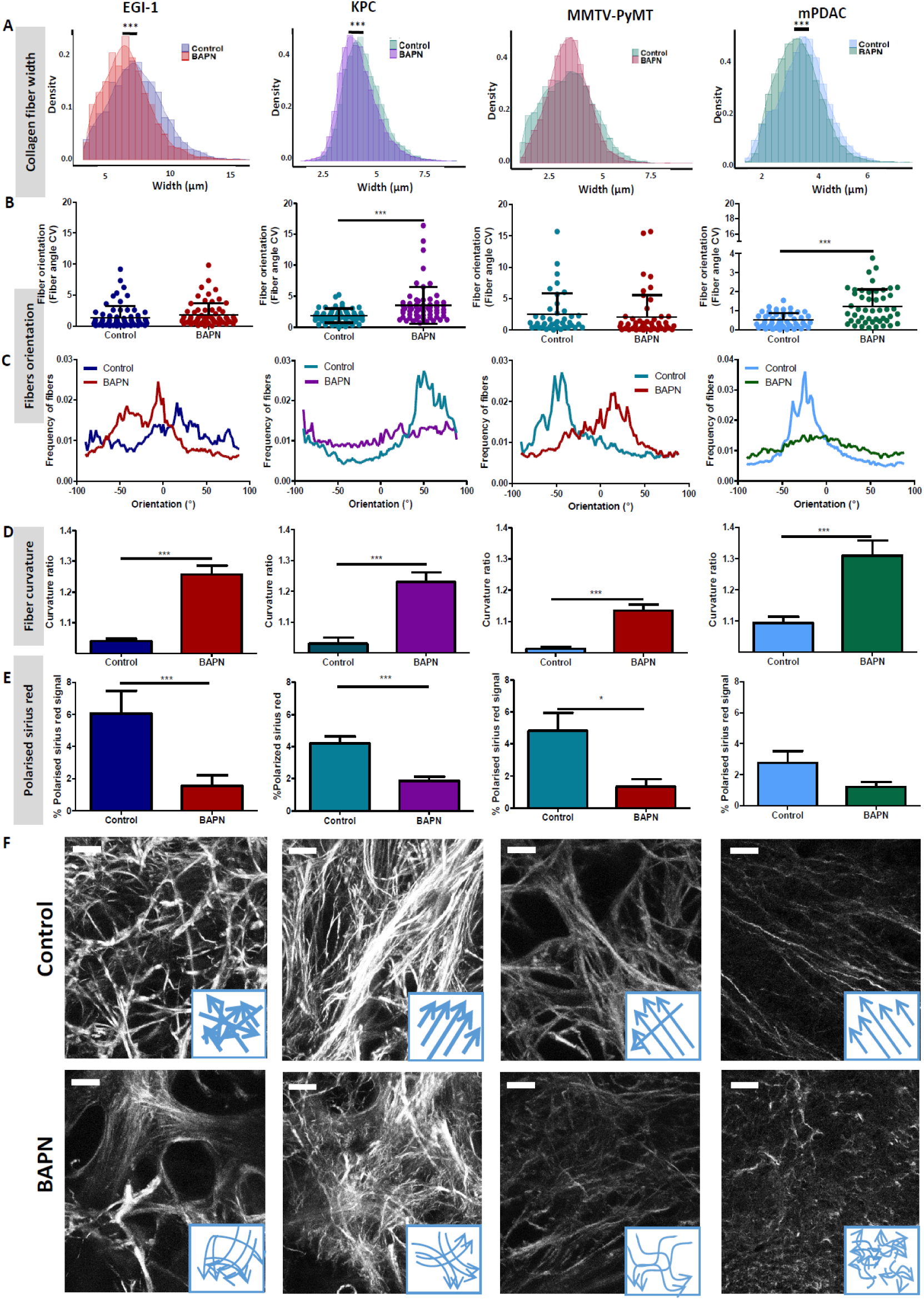
Effect of LOX inhibition on ECM architecture in EGI-1, KPC, MMTV-PyMT and mPDAC tumor models. (**A**) Collagen fiber width distribution measured from SHG images (\*\*\**P-*value< 0.001, Students t test, 50-70 images/ tumor, n=3). (**B**) Fiber orientation defined by the coefficient of variation (CV) (\*\*\**P-*value < 0.001, Students t test, 40-50 images/ tumor, n=3. (**C**) Representative example of fibers orientation distribution. (**D**) Collagen fiber curvature defined by the curvature ratio (\*\*\**P-*value< 0.001, Students t test, 40-50 images/ tumor, n=3). (**E**) Percentage of red-orange birefringent fibers combining red sirius staining and polarized microscopy, orange-red fibers correspond to thick and packed regions (\**P-*value<0.05, \*\*\**P-*value<0.001, Students’ t test, 20-30 images/ tumor, n=4). (**F**) SHG images of collagen networks in EGI-1, MMTV-PyMT, mPDAC and KPC control and BAPN-treated tumors. Illustrative scheme indicating the changes in width, orientation and curvature of collagen fibers induced by BAPN treatment. Scale bar = 50 µm.

A significant reduction of collagen fiber width distribution was observed in EGI-1, KPC and mPDAC models, whilst fibers in the MMTV-PyMT tumor model did not display a significant change in their width (**Figure 3A**). The most substantial differences was seen in the EGI-1 model, where collagen fiber width was decreased by 9,4 % on average (6.7µm versus 7.4 µm). Changes in mPDAC and KPC were less pronounced, with a reduction of 4.4 and 5 % respectively (**Table S2**). The inhibition of LOX did not affect collagen fiber length in any of the models (**Table S2**).

We next assessed the orientation and linearization of collagen fibers in control and BAPN-treated animals. In general, the collagen fibers in normal tissues are typically curly and anisotropic in contrast to the situation observed in progressing tumors in which many of the fibers progressively thicken and linearize. Collagen fiber orientation was described as the coefficient of variation (CV) of the angle for all fibers, the smaller the CV is, the more aligned the fibers are. Fibers in non-treated tumors remained mainly oriented in one dominant direction (**Figures 3B, 3C and 3F**) with a CV from 1.85 to 0.5 consistent with previous findings (*35*). LOX inhibition tends to disrupt the alignment of collagen fibers, meaning that they were more dispersed and oriented in different directions with increased CV as compared to control conditions. The most significant effects were seen in KPC and mPDAC model.

Collagen in tumors are characterized for being linear and reticulated due to the high level of deposition and posttranslational crosslinking. This physical restructuration of collagen progressively stiffness the ECM (*36*). The level of collagen linearization was quantified by measuring the curvature ratio of the fibers. The curvature ratio of control tumors for all models was close to 1 meaning that the collagen fibers were fully linearized. In contrast, LOX inhibition severely affected fiber curvature, as there was a reversion of the fibers linearization resulting in less linear and wavier fibers in all models (**Figures 3D and 3F)**.

Finally, LOX inhibition also significantly decreased the surface covered by thick and densely packed collagen fibers in EGI-1, KPC and MMTV-PyMT model (**Figures 3E and S6**), correlating with the significant decrease in average tumor stiffness and in the proportion of stiff areas measured *in vivo. In vivo* stiffness imaging and *ex vivo* characterization of the ECM structure were completed by *ex vivo* mechanical evaluation of isolated tissue samples with atomic force microscopy (AFM) nanomechanical measurements and plate shear rheometry at the tissue level in the KPC model. **Figure 4** shows the mean values of storage modulus (G’) of control and BAPN treated tumors. It can be noted that tumors treated with BAPN were softer than control ones, with mean stiffness of 1.80 ± 0.51 kPa compared 4.15 ± 1.92 kPa for the control. These results are in line with SWE measurements *in vivo* confirming a significant reduction in the mean stiffness of KPC tumors treated with LOX inhibitor. In contrast to bulk rheometry, AFM reveals the spatial heterogeneity of Young’s moduli at the sub-cellular level measured on 6 control (**Figure S7**) and 6 BAPN-treated tumors (**Figure S8**). For each sample high heterogeneity in tumors stiffness can be observed, as previously seen in other types of solid tumors like breast cancer (*37*). AFM measurements indicated that treatment of mice with BAPN leads to a narrower Young’s modulus distributions shifted to lower values of elastic modulus in comparison to the control samples (mean Young’s modulus of 0.82 ± 1.58 kPa versus 1.70 ± 1.66 kPa, respectively) (**Figures 4 B**). This confirms that the high heterogeneity in local tissues mechanical properties of KPC tumor can be reduced upon treatment with LOX inhibitor. Both local and global measurements confirm the normalization of tumor tissue mechanical properties mediated by LOX inhibition with drastic reduction in the linearized tightly packed collagen fibers that contribute to tumor stiffness heterogeneity and global enhancement in non-treated KPC tumors.

**Figure 4.**
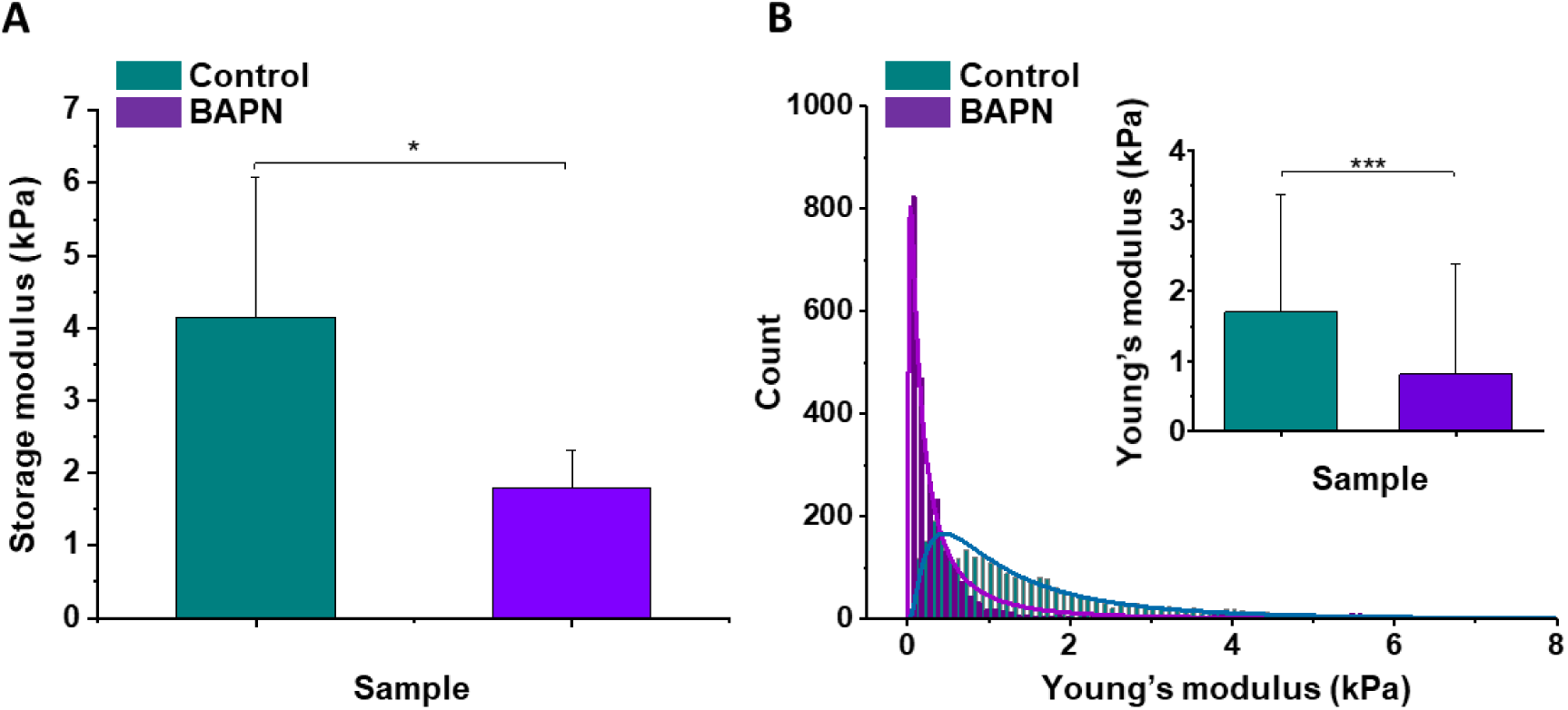
Mechanical properties of the mouse pancreatic ductal adenocarcinoma KPC tumor model and effects of LOX inhibition. (**A**) Rheological properties of tumor samples measured using plate shear rheometer. Mean storage modulus (G’) for all control and BAPN treated samples ± SD are presented (for 5% sample compression). (**B**) The Young’s modulus values distributions obtained for all control and treated tumor samples using AFM indentation technique. Inset in Figure 4B shows tissues’ Young’s modulus mean values ± standard deviation to highlight the difference between control and BAPN tissues. Statistical significance was determined using two-tailed Student’s t-test for overall values (n=6).

Overall, our results demonstrate the rationale of targeting LOX enzymatic activity for normalizing tumor mechanical properties and ECM structure (mostly collagen fibers compaction, segregation and linearity) in tumors exhibiting high tumor stiffness together with mechanical and structural heterogeneity.

### LOX inhibition increased intratumoral T cell migration and infiltration

Previous studies performed in our group have proved that the density and orientation of the ECM can have important impact on T cell behavior and their displacement in fresh human lung and ovarian tumor slices (*11, 38*). Motile T cells were mainly found in loose ECM stromal regions whereas fibrotic areas were devoid of lymphocytes. Based on this, we hypothesized that LOX-dependent tumor stiffness could consequently affect T cell migration in tumors and eventually predict the T cell behavior in the various ECM environment. To test this, we performed dynamic imaging of T cell migration on fresh tumor slices from mice treated or not with BAPN, using an approach previously established (*39*). EGI-1 is a xenografted tumor model, with implantation of human carcinoma cells into immune-suppressed mice that lack resident T cells. Thus, in order to evaluate T cell migration in this model we isolated human peripheral blood T cells (PBT) and activated them *in vitro*. We then added the activated PBT onto fresh tumor slices and analyzed their migration using real-time confocal microscopy. As the MMTV-PyMT tumor model is poorly infiltrated in host T cells, we investigated the migration of exogenously purified murine activated PBTs in the same manner as for the human EGI-1 model. In both mPDAC and KPC mice tumor models, resident tumor infiltrating T lymphocytes (TILs) were monitored after staining with directly-coupled anti-CD8 antibodies (*39*). The three parameters analyzed to assess T cell migration were cell migration speed (mean cell migration speed over 20 min), cell displacement (displacement vector between starting and final position) and straightness (ratio of cell displacement to the total length of the trajectory) of the migration trajectory (**Table 2, Figure S9**).

**Table 2.** T cell migration in the different tumor models and effect of LOX inhibition. Migration of activated PBT overlaid onto fresh tumor slices was analyzed in EGI-1 and MMTV-PyMT model, whilst resident tumor infiltrated T lymphocyte migration was analyzed in mPDAC and KPC models (* *P-* value<0.05, \*\*\**P-*value<0.001, Students’ t test, n=3-12 mice/group). T cell infiltration (CD8/mm^2^) calculated from immunofluorescence images. Results are shown as mean ± SD. Illustrative images of T cell migration tracks in EGI-1 tumor model. Tumor stroma (fibronectin) in red, tumor cells (EpCAM) in blue and T cells (Calcein) in green. Tracks are color coded to illustrate track displacement. Scale bar = 100 µm. See also **Movies S1 and S2**. TILs: Tumor infiltrating T lymphocytes.

| Tumor model | LOX inhibition | Type of T cell analyzed | T cell migration parameters |  |  | CD8/mm<sup>2</sup> |
| --- | --- | --- | --- | --- | --- | --- |
|  |  |  | Migration speed (μm/min) | T cell displacement (μm) | Track straightness (unitless) |  |
| EGI-1 | - | PBT deposit | 2.3 ± 1.5 | 12.11 ± 8.40 | 0.39 ± 0.20 | - |
|  | + |  | 1.9 ± 1.1 | **15.38 ± 10.30 \*\*\*** | **0.55 ± 0.20 \*\*\*** | - |
| KPC | - | TILs | 3.5 ± 2.1 | 15.09 ± 13.68 | 0.40 ± 0.25 | 252 ± 191 |
|  | + |  | **4.1 ± 2.7 \*** | 15.89 ± 12.11 | 0.40 ± 0.24 | **823 ± 591\*\*\*** |
| MMTV-PyMT | - | PBT deposit | 4.1 ± 2.2 | 13.15 ± 10.80 | 0.40 ± 0.24 | - |
|  | + |  | 3.9 ± 2.3 | **17.20 ± 15.90 \*\*\*** | 0.40 ± 0.25 | - |
| mPDAC | - | TILs | 1.0 ± 0.5 | 2.03 ± 1.40 | 0.10 ± 0.10 | 15.7 ± 10.0 |
|  | + |  | **1.6 ± 0.8 \*** | **8.80 ± 6.20 \*\*\*** | **0.50 ± 0.26 \*** | 21.2 ± 17.0 |

In control untreated tumors, T cells migrated slowly with average velocities which were relatively homogeneous in the different models except in the mPDAC model. Average velocities ranged from 2.3 µm/min in the EGI-1 model to 4.1 µm/min in the MMTV-PyMT model. These values are in line with a number of studies including ours showing a poor migration of T cells in tumors as compared to other organs (e.g. in lymph nodes) where T cell actively migrate (*40*). For example, the mean velocity of CD8 T cells in human lung tumors approaches 3 µm/min (*40*). Analysis of T cell track straightness gives indices close to 0.4 consistent with previous reports on T cell displacements in tumors. The mPDAC model differs from the others since T cells were almost static during the 20 min recording (average speed of 1 µm/min and straightness of 0.1).

In every tested model, LOX inhibition resulted in an overall increase in T cell migration as compared to control condition. However, different parameters were altered in each model depending on the nature of T cells that were monitored. In EGI-1 BAPN-treated tumors, activated PBT cells displayed longer displacement lengths compared to untreated tumors. This was also true for MMTV-PyMT tumors. Since activated PBT are not specific to the tumor, the effects observed are due to LOX inhibition and not due to T cells engaging in stable conjugates with cancer cells trough antigen recognition. In EGI-1 tumors, the trajectory straightness of activated PBTs was also significantly increased upon LOX inhibition.

When we evaluated the effect of BAPN on the dynamics of endogenous T cells infiltrated into KPC and mPDAC tumors, the most striking difference was found in the mPDAC model with a 5-fold increase in the displacement of T cells. In terms of cell speed, enhancements were observed in both the KPC and mPDAC models. The trajectory straightness was increased in mPDAC models but not in KPC tumors. In addition, we also evidenced an increased infiltration of resident T cell in tumors when treated with BAPN. Under anti-LOX treatment, a significant 4-fold increase of CD8+ T cell infiltration in both the stroma and tumor islets was observed in the KPC model (**Table 2, Figure S10**).

Results reported in **Table 2** have been obtained with data pooled from all mice either treated or not with BAPN. We decided to extend our analysis at the level of individual mouse and investigated the relationship between T cell motility (speed and displacement) and mean stiffness of control and BAPN-treated tumors. Our data indicate that T cell motility was inversely correlated to tumor stiffness (**Figure 5**) in line with Table 2 and Figure S9. However, in three out of the four models tested this correlation is different if one compares control and BAPN-treated tumors. In BAPN-treated tumors, there is a clear inverse linear correlation between T cell migration speed and mean tumor stiffness as evidenced by a steep slope. In comparison, such correlation is less pronounced in control stiff tumors. Similar results were observed when comparing T cell displacement and mean tumor stiffness. These results suggest that above a stiffness threshold that depends on the model, T cells are mostly arrested. In contrast, as tumor stiffness decreased by inhibition of LOX, T cell migration was restored. Thus, T cell motility is highly influenced by small variations in the stiffness of softer tumors as it is the case when LOX’s activity is inhibited.

**Figure 5.**
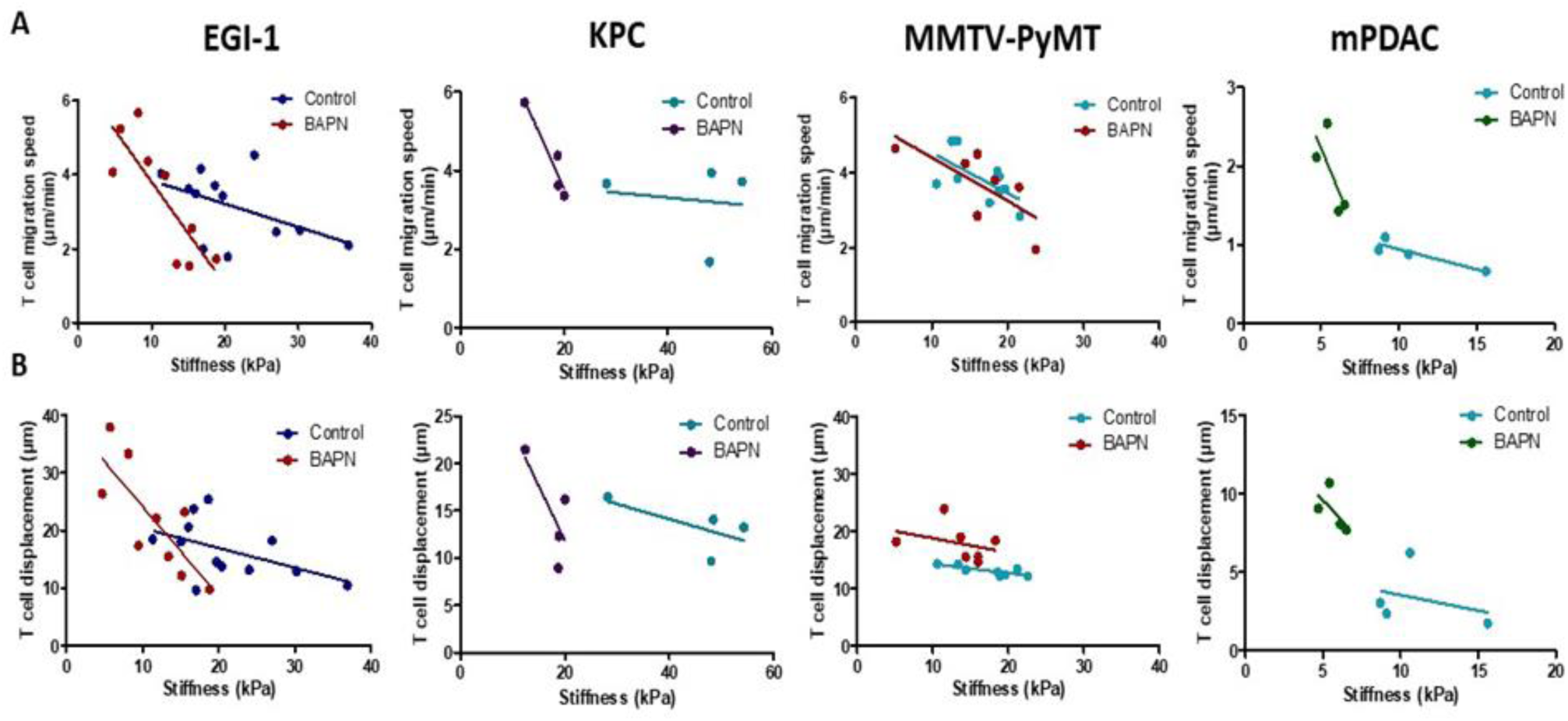
Correlation between mean tumor stiffness and T cell migration parameters in EGI-1, KPC, MMTV-PyMT and mPDAC tumor models. (**A**) Correlation between mean tumor stiffness and mean T cell velocity. (**B**) Correlation between mean tumor stiffness and mean T cell displacement. Averaged T cell velocity and displacement were calculated from at least 50 individual cells. Each point represents values from an individual mouse.

Overall, these results suggest that the excessive accumulation and linearization of collagen in ECM limits T cell migration within several rigid tumors with desmoplastic evolution and that LOX-inhibiting BAPN treatment can both reverse tumor stiffening and improve T cell infiltration and migration to tumor cells. We also identify SWE tissue stiffness as a predictive physical marker of T cell motility and infiltration in desmoplastic tumors.

### LOX inhibition improves response to anti-PD-1 therapy

Even though the inhibition of LOX was followed by an increase in CD8 T cell number and migration, this finding was not accompanied with major effects on tumor growth in 4 out of the 5 tumor models tested (**Figures S1-S5**). In different settings, an increase in intratumoral T cell motility is not sufficient to reduce tumor growth if T cells are still impaired in their capacity to respond to tumor antigens (*39*). Consequently, we decided to assess whether LOX inhibition could improve the response to immune checkpoint inhibitors. KPC tumor bearing mice were treated with BAPN combined with anti-PD-1 antibodies. Mice were treated or not with BAPN from tumor cell injection up to their sacrifice and were treated with anti-PD-1 antibodies when the tumor volume was around 80 – 150 mm^3^. At this point the mice received 4 doses i.p. injection of anti–PD-1 or isotype control antibody at 4 days intervals (i.p. injection). As shown in **Figure 6A**, while the anti-PD-1 alone was not able to reduce tumor growth, the combination of BAPN with the checkpoint inhibitor significantly delays tumor progression. We then profiled the immune cell population in these tumors. We found that BAPN treatment alone significantly decreases the number of polymorphonuclear neutrophils (**Figure 6B**), but increases the presence of MHCII+ tumor associated macrophages (TAMs) (**Figure 6C**) while the combination therapy expanded the percentage of GrzmB CD8+ T cells (**Figure 6D**) and the ratio of CD8+ to Treg cells (**Figure 6E**). We also analyzed the amount of cytokines in supernatants of whole-tumor slices derived from these experiments. Results show that the combination therapy led to an increase in TNFα and RANTES, supporting further the increase of T cell infiltration and activation in this condition. In both BAPN and BAPN combined with anti-PD-1 conditions we observed an increase of GrzmB+ levels compared to the control condition.

**Figure 6.**
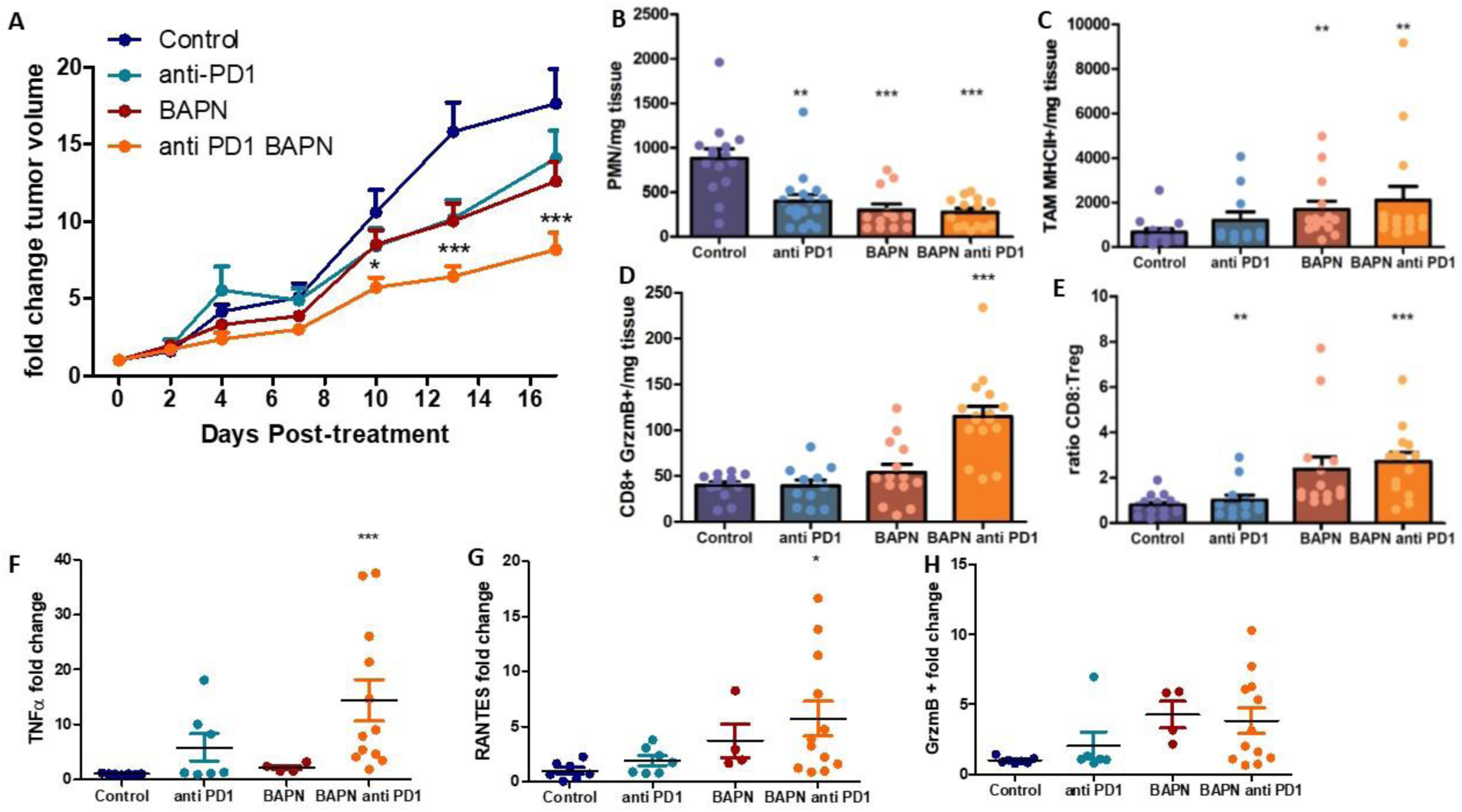
LOX inhibition increases the efficiency of anti-PD-1 therapy in KPC tumor model. (**A**) Tumor volume fold change after combination therapy of BAPN with anti-PD-1. (**B**) CD11b+, Ly6G+, Ly6C+ PMN/mg of tissue. (**C**) CD64+F4/80+MHCII+ TAMs/mg of tissue. (**D**) CD8+ T cells/mg tissue. (**E**) CD8+ to FoxP3+ Treg ratio. (**F-H**) Multiplex analysis of inflammatory chemokines (TNFα, RANTES and GrzmB) produced by fresh KPC tumor slices kept in culture for 18 h (*** *P-*value < 0.001, * *P-* value < 0.05, One-Way ANOVA, Krustal Walis, n= 8 per condition, two independent experiments).

Overall, while ECM and stiffness normalization achieved though LOX inhibition increases T cell infiltration and migration, this strategy also improves the efficacy of anti-PD-1 blockade

## DISCUSSION

Despite the success of targeting the stromal compartment in tumors (*7, 11, 15, 19, 20, 41*), in particular tumor ECM, there are still a series of challenges that remained to be addressed. In the first part of this study we tackle two of these challenges. One of them is finding ways to accurately assessing the architecture of the stroma. In this paper, we propose a thorough analysis combining non-invasive imaging techniques for a macroscopic characterization of tumor stiffness with advanced microscopy techniques to elucidate the collagen network structure, one of the most important components of the tumor ECM. We established a link between the architecture of the collagen network in different desmoplastic tumors with the tumor mechanical properties. This allowed us to extract mechanical and structural reference values and allowed to later evaluate the effect of an ECM targeted therapy. Such an approach could be translated to patients given the availability of SWE techniques that could be correlated to histological observations of the TME.

Accurately modeling tumor complexity and heterogeneity found in patients’ tumors using a single preclinical model, is a real challenge. In the first place, many preclinical models do not always reproduce the carcinoma structure found in their respective human tumors. Another important feature of human solid tumors, often absent in mouse tumor models, is their considerable higher stiffness compared to their neighboring healthy tissues which is highly correlated with cancer progression and metastasis. These two features can often be found in human xenograft models that, on the other hand, are not suitable to study immune reactions. For all of these reasons, in this study we have set up and characterized five different solid tumor models covering 3 types of carcinoma, in order to be able to tackle the different aspects of the TME as well as cover the heterogeneity found in patients.

Once the tumor structure and mechanical properties of each model were characterized, an ECM-targeting therapy dependent on LOX-inhibition with BAPN was tested. Previous studies performed by Levantal *et al*. in a spontaneous mouse model of breast carcinoma, showed that LOX inhibition induces structural changes in the collagen network (*31*). Here, we confirm and extend these results in other models and provide multiscale insights about different effects of LOX-inhibition from model to model, including stiffness measurements at the tissue level. Whilst for cholangiocarcinoma (EGI-1) and pancreatic adenocarcinoma models (KPC and mPDAC), LOX-inhibition drastically decreases tumor stiffness, this effect is less marked in the breast adenocarcinoma model (MMTV-PyMT). Collagen fiber curvature is affected in all models, however changes in fiber orientation are only significant in KPC and mPDAC models. These changes can be explained by the fact that the tumor microenvironment and thus the basal collagen structure in each tumor model is different, hence the inhibition of collagen cross-linking modifies tumor collagen network to different extents. This emphasizes the need of developing reliable diagnostic markers, such as SWE stiffness mapping, based on a clear understanding of the tumor collagen network, in order to predict the response to ECM targeting strategies.

It is well established that the number of T cells found within a tumor as well as their ability to migrate and reach cancer cells is key for an effective antitumoral response. These last few years a lot of efforts have been made in identifying cells and factors controlling the migration of T cells within tumors. The notion that prevailed is that growing tumors are composed of cells and factors, such as macrophages and hypoxia, hostile for T cells to migrate (*40, 42*). The importance of the ECM in controlling the distribution and migration of T cells in tumors has also emerged. In human lung and ovarian tumors we found that T cells preferentially accumulate and migrate in stromal regions that exhibited a loose matrix architecture but not in dense regions (*11, 38*). Likewise, in aged skin, dermal fibroblasts harbor a phenotype similar to CAFs and produce ECM matrices that limit T cell displacements (*43*). However, in triple-negative breast cancers and in pancreatic tumors T cells were still found in dense networks of collagen fibers (*35, 44*). Since most of the aforementioned studies were correlative, we decided in this study to specifically alter the ECM network by LOX inhibition and investigated the consequences on T cell motile behavior. Our results confirmed the importance of the ECM and tissue stiffness in controlling the migration of T cells in tumors. However, not all tumor models react similarly to BAPN-treatment and differences were observed on T cell motility parameters. In KPC and mPDAC models, LOX-inhibition results in an increase in TIL migration speed, whilst in the EGI-1 and MMTV-PyMT models the impact was mainly on T cell displacement. These differences in behavior can be explained by i) the fact that in the former case we analyzed resident TIL and in the later ex vivo purified and activated PBT that are not specific to the tumor and by ii) the fact that LOX inhibition causes differential effects in each of the models that can translate to differences in the impact on T cell migration. Regardless of the explanation, T cell intratumoral motility was inversely correlated to tumor stiffness as measured non-invasively using SWE (**Figure 5**) and this was true in all tested models. However, this relationship is not linear and in most models T cells strongly decelerate when a threshold in stiffness is reached. In soft tumors such as those induced by LOX inhibition, T cells manage to migrate. Conversely, in stiff non-treated control tumors T cell migration is impeded. These data fits well with results obtained *in vitro* in a range of 3D collagen matrices showing that T lymphocytes have the ability to adapt their morphology to the structure of the tissue up to a certain limit (*45*). In dense collagen matrices, T cell motility is halted. Our analysis supports the idea that elastography measurements could provide valuable companion markers to evaluate the need for anti-stromal strategy in order to normalize tumor stiffness and consequently improve T cell migration.

In this study we did not take into consideration possible effects of LOX inhibition on tumor blood vessels. Previous studies have reported a reduced angiogenesis after LOX and LOX-like protein inhibition and an increased perfused vessel density in the case of overexpression of LOX (*46, 47*). This could partly explain why we observe a significant increase in T cell infiltration in KPC tumors upon BAPN treatment. However, other recent studies argue the opposite as an increase in collagen crosslinking and matrix stiffness resulted in an increase in angiogenic sprouting. Conversely, the inhibition collagen cross-linking in tumors resulted in reduced vasculature density and permeability (*48*).

Given the low efficacy of T cell based immunotherapies in solid tumors any method to increase its effect on tumor regression is of interest. With the exception of desmoplastic melanomas, features of wound healing and fibrosis are usually detrimental to anti-PD-1 responses (*49, 50*). Accordingly, a number of anti-fibrotic strategies have been recently implemented in combination with immune checkpoint inhibitors (*10*). One of the most promising targets appears to be TGFβ. In preclinical mouse tumor models, TGFβ inhibition with immune checkpoint blockade induces complete and durable responses in otherwise unresponsive tumors. However, due to TGFβ pleiotropic effects, concerns regarding blockade of this cytokine arose (*7, 16*).

Our study indicates that LOX represents another valuable target as its inhibition in the transplanted KPC model increases the efficacy of anti-PD-1 treatment, while monotherapy with either agent alone is ineffective. Moreover, the combination treatment was associated with a tumor microenvironment shifted towards antitumoral effector cells and components whereas immunosuppressive cells were reduced. The reason of this reprogramming is not known for the moment. Apart from T cells, other immune cells can be subjected to regulation by the ECM. For instance, recent crosstalk between tumor-associated macrophages and ECM have been reported (*51*).

Although the clinical use of BAPN has been impeded by concerns regarding toxicities, other strategies to inhibit LOX in cancer and fibrotic disease patients are currently ongoing (*52*). Our work confirms LOX as a molecular target to improve T cell migration dynamics as well as to ameliorate the immunosuppressive microenvironment. It paves the way for clinical trials combining LOX inhibitors with PD-1/ PD-L1 blockade, possibly in biomarker-selected cohorts of patients with high tumor stiffness evaluated with non-invasive imaging approaches.

## MATERIALS AND METHODS

### Cell culture

Murine cell lines KPC (obtained from Corinne Bousquet – Université Toulouse III), MET-1 cell line (obtained from Robert Cardiff - University of California, Davis) and mPDAC (obtained from Douglas Hanahan -Swiss Institute for Experimental Cancer Research) were cultured in DMEM supplemented with 10 % Fetal bovine serum (FBS), Penicillin/Streptomycin, L-Glucose and Sodium pyruvate (Gibco). EGI-1 cells, derived from extrahepatic biliary tract, were obtained from the German Collection of Microorganisms and Cell Cultures (DSMZ,Germany), were cultured in DMEM supplemented with 1 g/L glucose, 10 mmol/L HEPES, 10% fetal bovine serum (FBS), antibiotics (100 UI/mL penicillin and 100 μg/mL streptomycin) and antimycotic (0.25 μg/mL amphotericin B) (Invitrogen).

Human and mouse CD8+ T cells were isolated and cultured as described in the Supplemental experimental procedures.

### In vivo studies

All animal experiments were performed in agreement with institutional animal use and care regulations after approval by the animal experimentation ethics committee of Paris Descartes University (CEEA 34, 16-063). For details on protocols used for every mouse models, see Supplemental methods.

Tumor growth was followed with a caliper, and tumor volume (V) was calculated as follows: xenograft volume = (xy^2^)/2 where x is the longest and y, the shortest of two perpendicular diameters.

For LOX inhibition, animals were treated with BAPN (3mg/mL, Sigma) in the drinking water, which was changed twice per week. In all implantable models (KPC, mPDAC, MET-1 and EGI-1) animals were treated from the day of the tumor implantation until the end of the study. For the MMTV-PyMT model, animals were treated at 10-week of age, approximately the moment tumor start developing.

Experiments combining LOX inhibition and anti-PD1 were conducted in the KPC model. Four i.p. injections of 200 µg of anti–PD-1 antibody (RMP1-14 clone; BioXcell) or isotype control (rat IgG2a; BioXCell) were started when tumors reached 80 −100 mm^3^ size. Injections were performed every 4 days.

### Shear wave elastography

Shear wave elastography (SWE) measurements were performed every 3-4 days during the entire follow-up of the tumor growth. Images were acquired with the ultrasound device Aixplorer (SuperSonic Imagine, Aix-en-Provence, France) using a 15-MHz superficial probe dedicated to research (256 elements, 0.125 µm pitch). See Supplemental information for more details.

### Tumor slice imaging

Tumor slices were prepared following the protocol described elsewhere (*12, 38, 39*). Briefly, samples were embedded in 5% low-gelling-temperature agarose (type VII-A; Sigma-Aldrich) prepared in PBS. Slices (350 μm) were cut with a vibratome (VT 1000S; Leica) in a bath of ice-cold PBS.

In EG-1 and MMTV-PyMT models, T cell migration was assessed on previously purified and activated T cells introduced into fresh tumor slices. In KPC and mPDAC models, resident CD8 T cell migration was evaluated after staining for 15 minutes at 37°C with PerCP-e710 anti-mouse CD8a (53-6.7 clone, eBioscience). For details see Supplemental Information.

### Dynamic imaging analysis

A 3D image analysis was performed on x, y, and z planes using Imaris 7.4 (Bitplane AG). First, superficial planes from the top of the slice to 15 μm in depth were removed to exclude T cells located near the cut surface. Cellular motility parameters were then calculated. Tracks >10% of the total recording time were included in the analysis.

### Second harmonic generation microscopy

The images were obtained using an inverted stand Leica SP5 microscope (Leica Microsystems GmbH, Wetzlar, Germany) coupled to a femtosecond Ti:sapphire laser (Chameleon, Coherent, Saclay, France) tuned at a wavelength of 810 or 850 nm for all experiments. See Supplemental information for more details.

### Histology

For most of the tumors, half of the biopsy was fixed overnight at 4 °C in a periodate–lysine– paraformaldehyde solution [0.05 M phosphate buffer containing 0.1 M L-lysine (pH 7.4), 2 mg/mL NaIO4, and 10 mg/mL paraformaldehyde]. See Supplemental information for more details.

### Atomic Force Microscopy and shear rheometry

Mechanical measurements of the control and BAPN mice tumors were performed *ex vivo* at the nanoscopic and macroscopic scale using atomic force microscopy (AFM) and shear rheometer.

Firstly, millimeter-scale samples of mice KPC model tumors were measured with a JPK Bruker NanoWizard 4 BioScience atomic force microscope working in the force spectroscopy mode. For the macroscopic rheological tests HAAKE Rheostress 6000 rheometer (Thermo Fisher Scientific, Waltham, MA, USA), fitted with 8 mm diameter parallel plate system was used. See Supplemental information for more details.

### Flow cytometry

Tumors were mechanically dissociated and digested for 45 min at 37 °C in RPMI 1640 with 37.5 µg/mL Liberase TM (Roche) and 8,000 U/mL DNase I, bovine pancreas (Merck Millipore). The resulting digestion was filtered through a 70 µm cell stainer and centrifuged. Red blood cell lysis with ACK buffer was performed on the remaining pellet and subsequently filtrated on a 40-µm cell strainer, the cell suspension was rinsed in PBS and stained in 96-well round-bottom plates with a LIVE/DEAD Fixable Blue Dead Cell Stain Kit (Invitrogen) for 20 min at 4 °C. Cells were then washed and stained with Abs for surface markers for 20 min at 4 °C. More details and the list of Abs are in Supplemental Information.

### Cytokine detection in tumor slice supernatants

Fresh KPC tumor slices were prepared as previously described and kept at 37 °C in 24-well plates with 0.5 mL RPMI per well. Four to five slices were put in culture for each mouse. Eighteen hours later supernatants were collected and centrifuged at 300 × g to eliminate suspension cells. Cell-free supernatants were frozen and stored at 80 °C. Granzyme B, TNFα and RANTES (CCL5) release was assayed by Luminex technology (Bio-Plex 200 from Bio-Rad) with a customized Milliplex kit (Merck Millipore).

### Statistical analysis

Results were analyzed using the GraphPad Prism 5.0 statistical software. Data are shown as means ± standard error of the mean (SEM). For comparisons between two groups, parametric Student t-test or non-parametric Mann-Whitney test were used. For comparisons between more than two groups, a parametric One-Way analysis of variance (ANOVA) test was followed by a posteriori Kruskal-Walis test.

## SUPPLEMENTAL INFORMATION

Supplemental information includes 10 figures, 2 movies, supplemental experimental procedures, 2 supplemental tables, one supplemental reference.

## ACKNOWLEDGMENTS

This study was supported by grants from the French Ligue Nationale contre le Cancer (Equipes labellisées) (ED), Plan Cancer (Tumor heterogeneity and ecosystem program) (ED), CARPEM (Cancer Research for Personalized Medicine) (ED). ANB received a PhD fellowship by the Institute thematique multi-organismes (ITMO) Cancer, the doctoral school Frontières du Vivant (FdV) – Programme Bettencourt and the Fondation ARC pour la recherche sur le cancer. LF and JV are member of the European Network for the Study of Cholangiocarcinoma (ENSCCA) and participates in the initiative COST action EURO-CHOLANGIO-NET granted by the COST Association (CA18122). We would like to thank the staff of the IMAG’IC, CYBIO, PIV and HistIM facilities of the Cochin Institute for their advice all along this study. We acknowledge Tatiana Ledent from Housing and experimental animal facility (HEAF), and Fatiha Merabtene and Brigitte Sohlonne from the histomorphology platform, Centre de Recherche Saint-Antoine (CRSA). Dr Joanna Mystkowska and Dawid Lysik (a PhD student) from the Department of Materials Engineering and Production, Faculty of Mechanical Engineering from Bialystok University of Technology for facilitating the measurements using the rheometer.

## Author Contributions

ANB, GR, IC, JC, LF, FG and ED conceived the project, designed and performed the experiment, analyzed the data and wrote the manuscript. JV, SB, CKM and MP contributed to the in vivo experimental section. PD, KP, RB performed AFM and rheological measurements. All authors discussed the results and commented the manuscript.

## SUPPLEMENTARY INFORMATION

### SUPPLEMENTAL DATA

#### Supplementary figures

**Figure S1.**
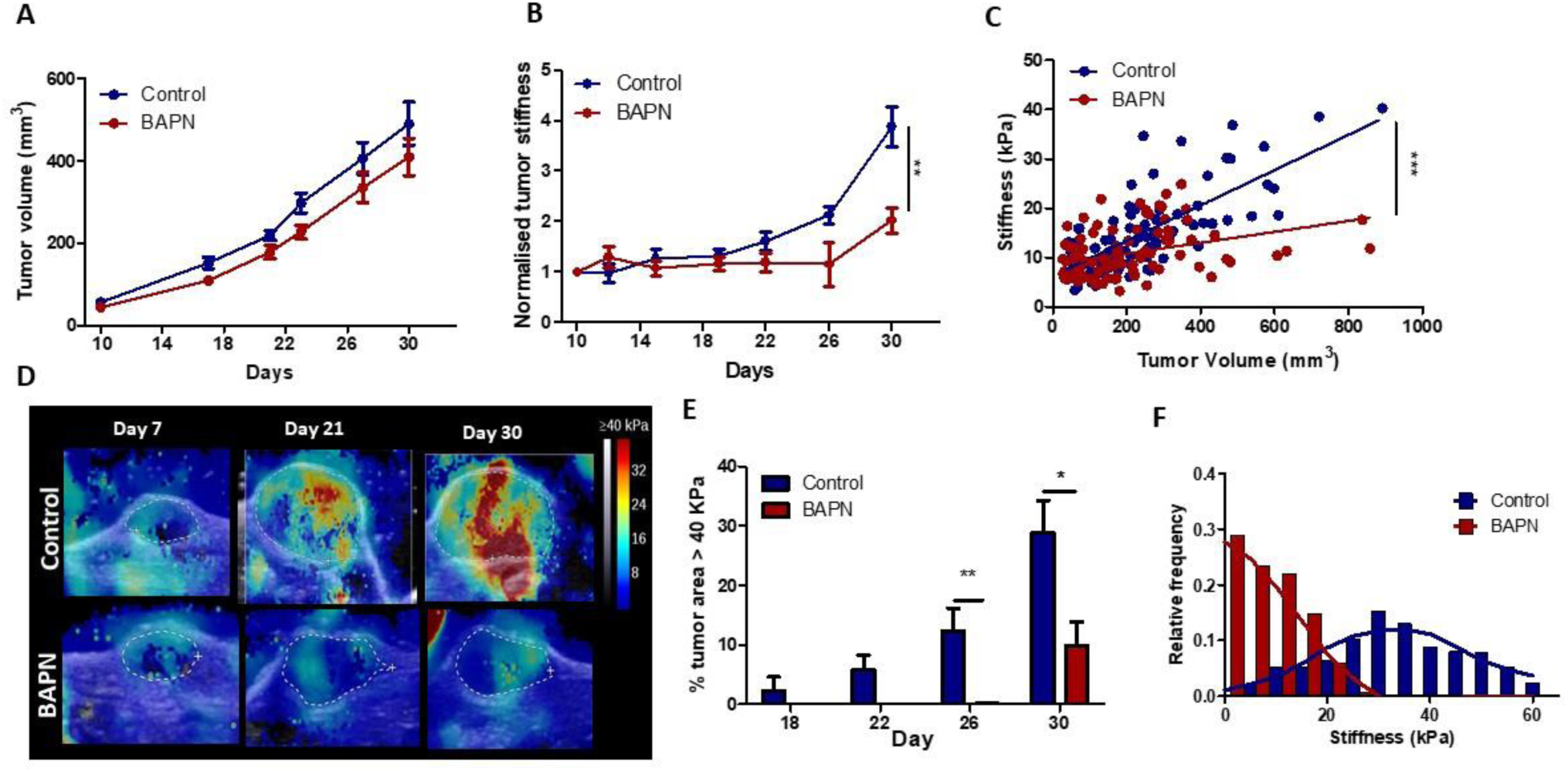
Effect of LOX inhibition in EGI-1 tumor model. (**A**) Evolution of tumor growth in control and treated tumors (n = 30/ group, three independent experiments). (**B**) Normalized tumor stiffness of EGI-1 control and BAPN treated tumors. (**P-value < 0.01, Students t test, n=30, three independent experiments). (**C**) Correlation of tumor mean stiffness and tumor volume for control and treated tumors. (***P-value < 0.01 ANCOVA). (**D**) Representative SWE images of control and BAPN-treated tumors at day 7, 21 and 30 of treatment. (**E**) Percentage of the tumor with stiffness > 40 kPa in tumors at day 18, 22, 26 and 30 of treatment. (**F**) Representative histogram of stiffness distribution in a control and a BAPN-treated tumor at day 30.

**Figure S2.**
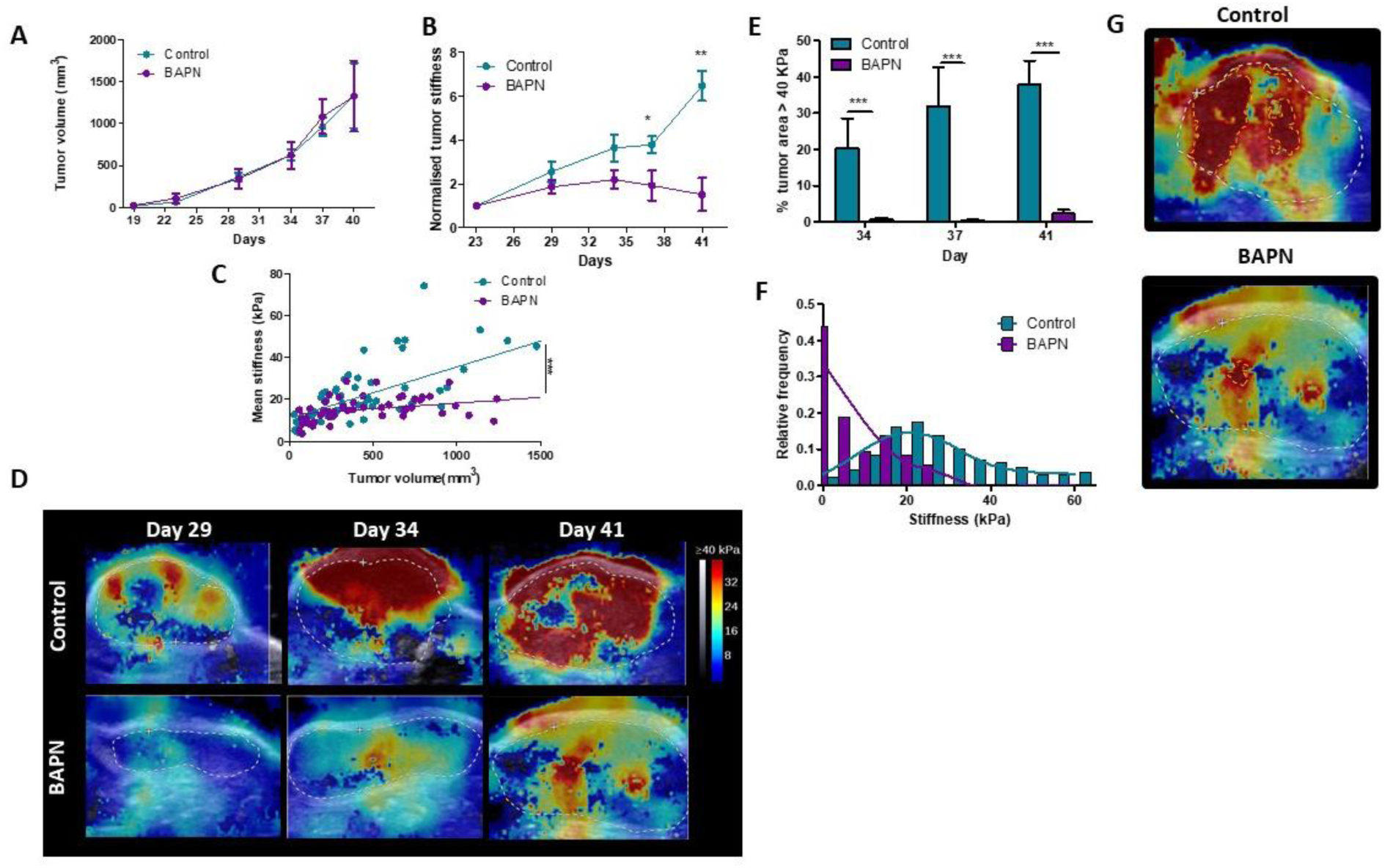
Effect of LOX inhibition in KPC tumor model. (**A**) Evolution of tumor growth in control and treated tumors (n = 34/ group, three independent experiments). (**B**) Normalized tumor stiffness of KPC control and BAPN treated tumors. (\**P-*value< 0.05, ** *P-*value < 0.01, Students t test, n=34, three independent experiments). (**C**) Correlation of tumor mean stiffness and tumor volume for control and treated tumors. (\*\*\**P-*value< 0.01 ANCOVA). (**D**) Representative SWE images of control and BAPN-treated tumors at day 29, 34 and 41 of treatment. (**E**) Percentage of the tumor with stiffness > 40 kPa in tumors at day 34, 37 and 41 of treatment. (**F**) Representative histogram of stiffness distribution in a control and a BAPN-treated tumor at day 41. (**G**) Representative SWE images of stiff regions (> 40kPa) in control and treated tumors

**Figure S3.**
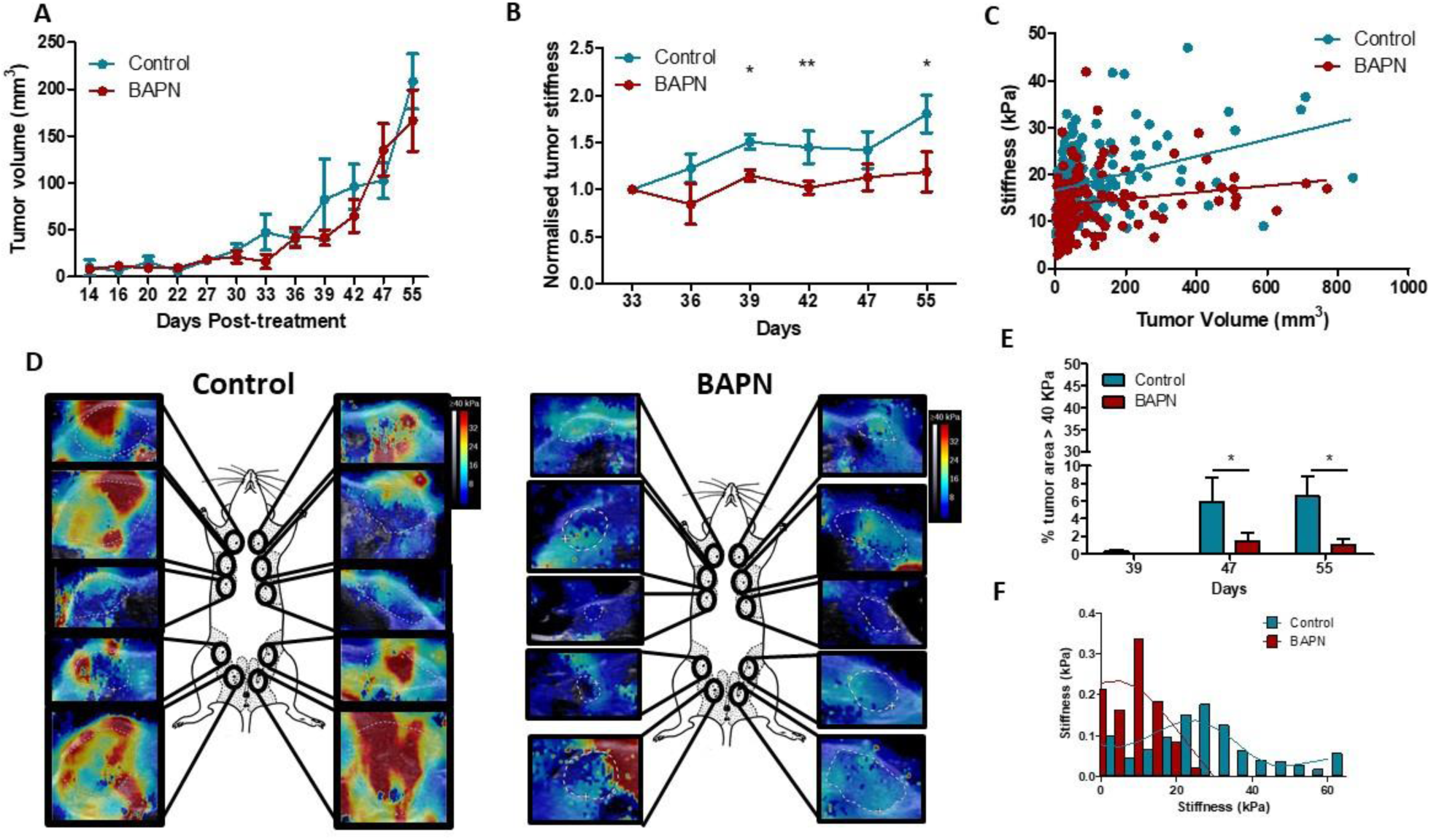
Effect of LOX inhibition in MMTV-PyMT tumor model. (**A**) Evolution of tumor growth in control and treated tumors (n = 5/ group, 10 tumors per mouse). (**B**) Normalized tumor stiffness of MMTV-PyMT control and BAPN treated tumors. (\**P-*value < 0.05, ** *P-*value < 0.01, Students t test, n=30, n = 5/ group, 10 tumors per mouse). (**C**) Correlation of tumor mean stiffness and tumor volume for control and treated tumors. (**D**) Representative SWE images of control and BAPN-treated tumors at day 42 of treatment. (**E**) Percentage of the tumor with stiffness > 40 kPa in tumors at day 39, 47 and 55 of treatment. (**F**) Representative histogram of stiffness distribution in a control and a BAPN-treated tumor at day 55.

**Figure S4.**
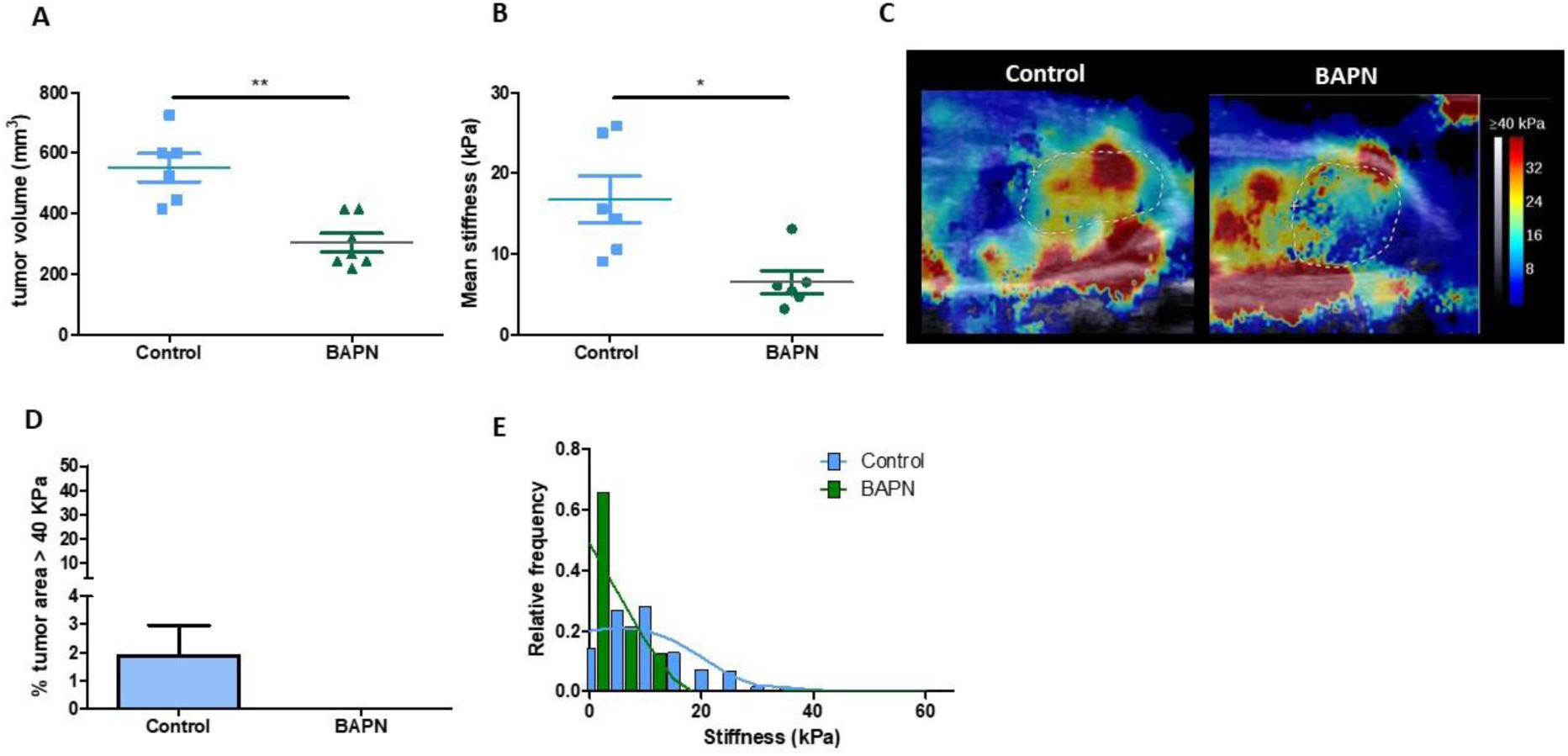
Effect of LOX inhibition in mPDAC tumor model. (**A**) Tumor volume measured by ultrasound imaging, at day 21 of treatment (\*\**P-*value < 0.01, Student t test, n = 7/ group). (**B**) Mean tumor stiffness of mPDAC control and BAPN treated tumors. (\**P-*value < 0.05, Student t test, n = 7/ group). (**C**) Representative SWE images of control and BAPN-treated tumors at day 21. (**D**) Percentage of the tumor with stiffness > 40 kPa in tumors at day 21. (**E**) Representative histogram of stiffness distribution in a control and a BAPN-treated tumor at day 21.

**Figure S5.**
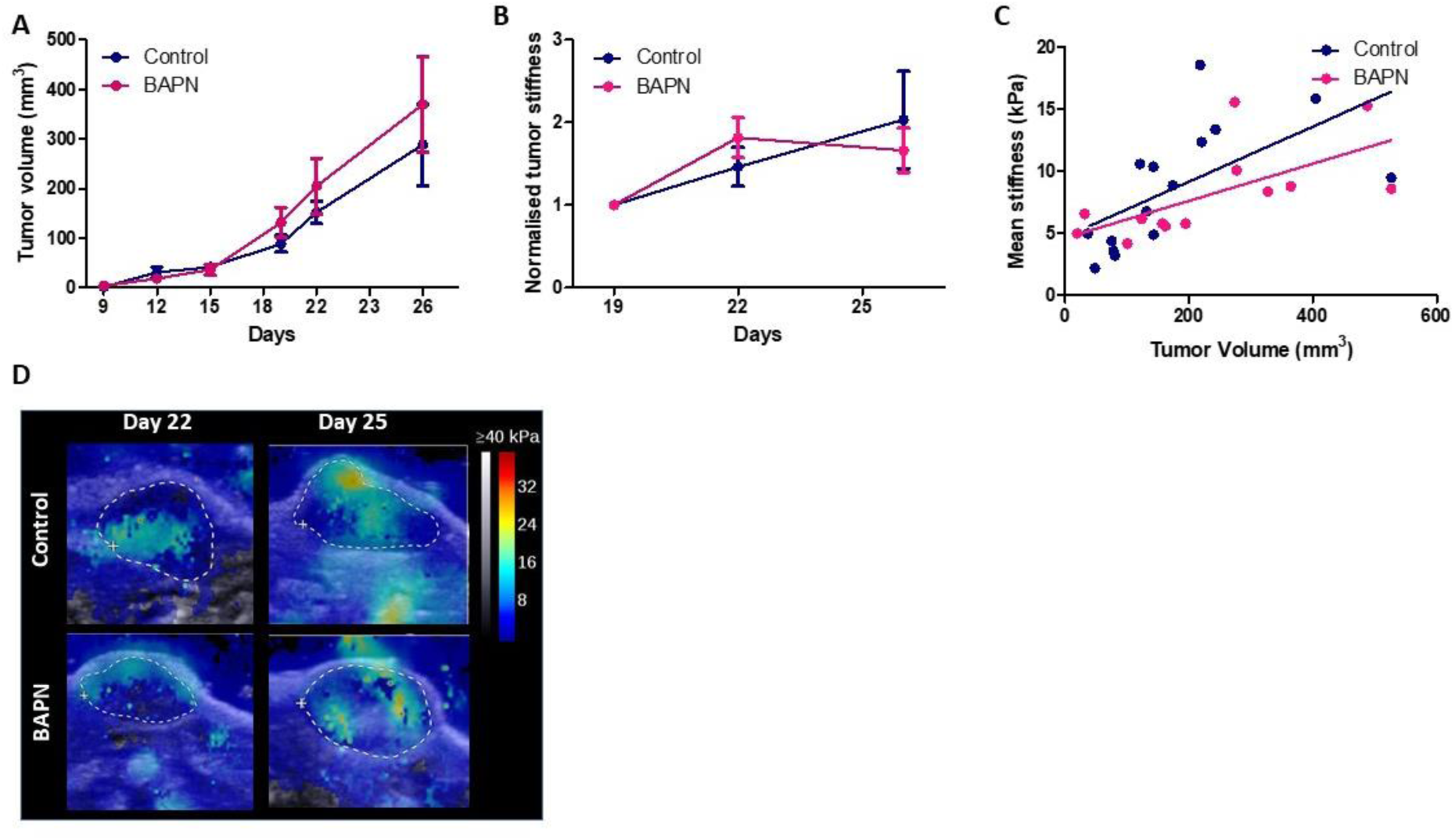
Effect of LOX inhibition in MET-1 tumor model. (**A**) Evolution of tumor growth in control and treated tumors (n = 12/ group). (**B**) Normalized tumor stiffness of MET-1 control and BAPN treated tumors (n = 12/ group). (**C**) Correlation of tumor mean stiffness and tumor volume for control and treated tumors. (**D**) Representative SWE images of control and BAPN-treated tumors at day 26 of treatment.

**Figure S6.**
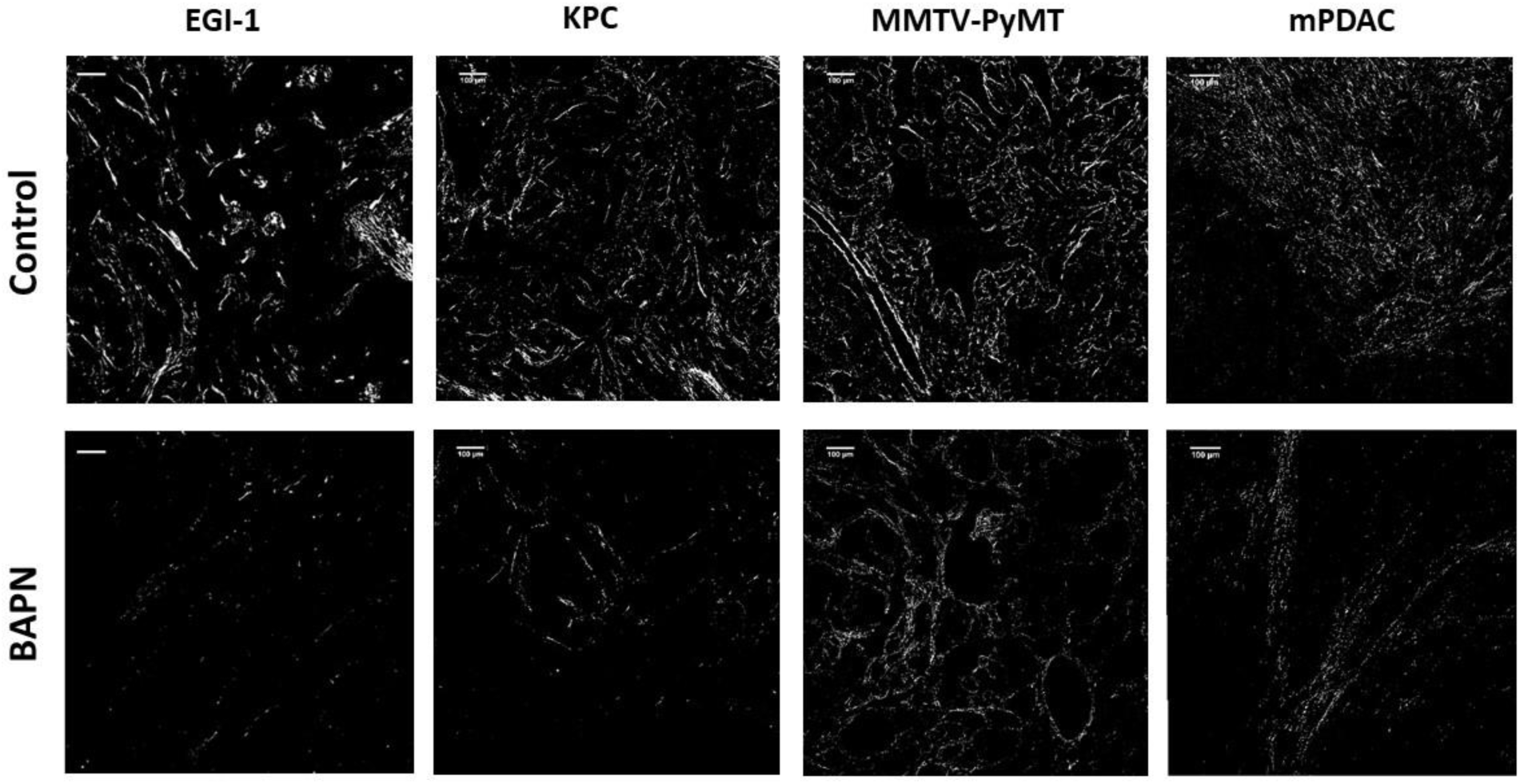
Combination of Sirius red staining and polarized microscopy. Surface covered by red-orange birefringent fibers that correspond to thick and packed regions in EGI-1 MMTV-PyMT, mPDAC and KPC tumor models. Scale-bar = 100 µm.

**Figure S7.**
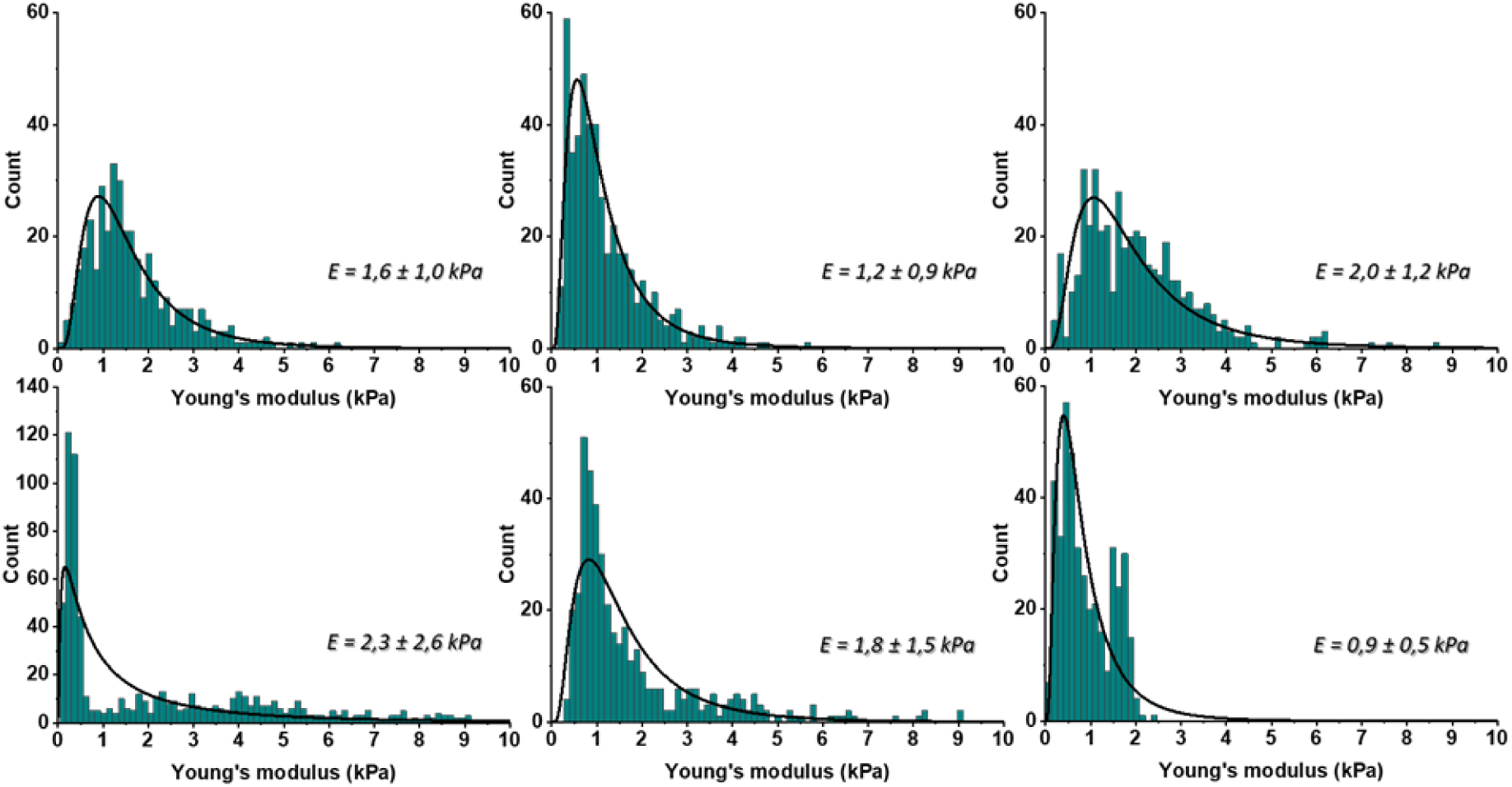
The distributions of Young’s moduli obtained for the control mouse pancreatic ductal adenocarcinoma KPC tumor model (Control) using AFM indentation technique with maximal indentation depth by AFM probe equal 2µm. Distributions were fitted with probability density function of the log normal distribution and corresponding mean ± SD values are presented on the right side of each distribution.

**Figure S8.**
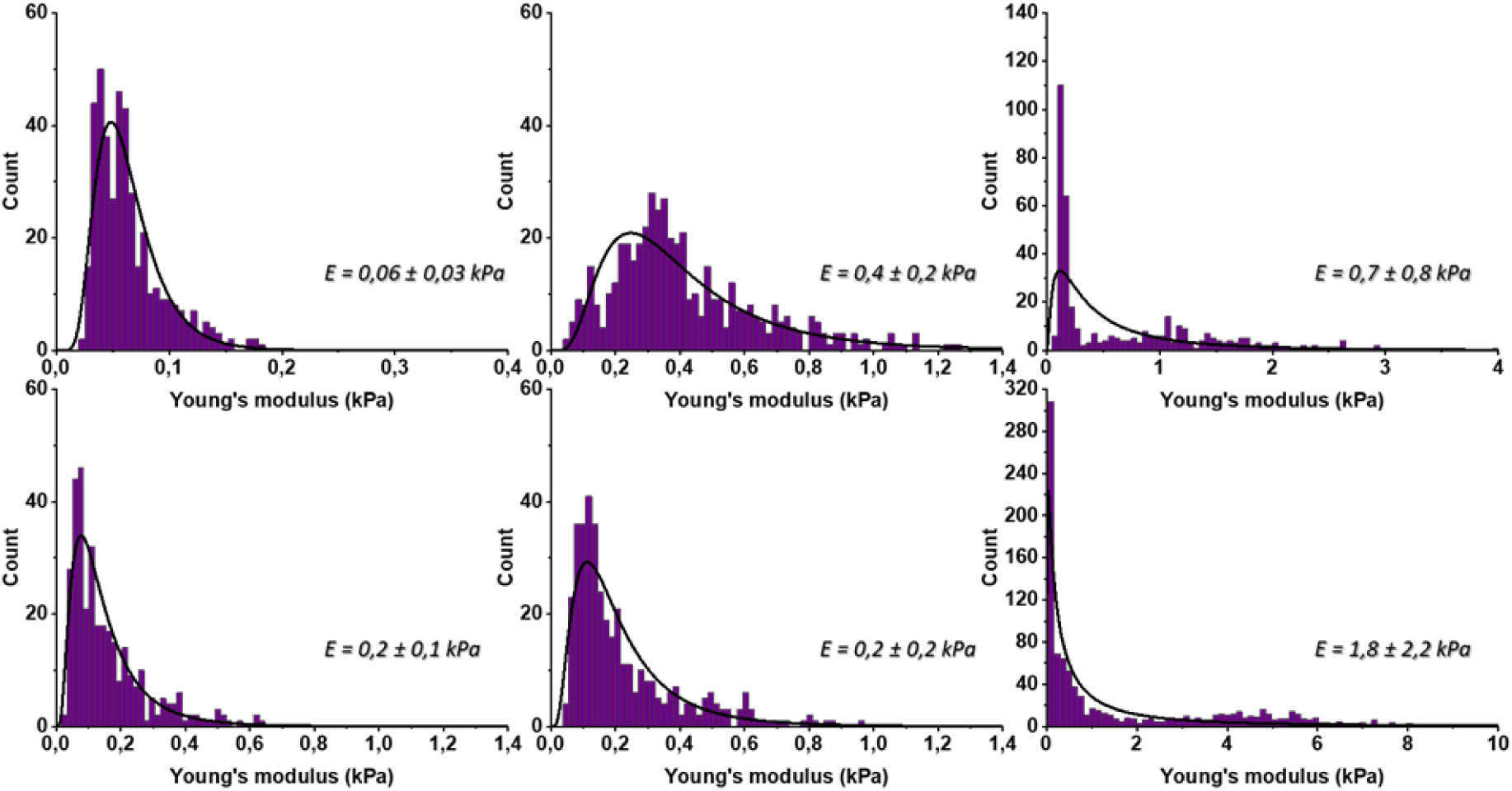
The distributions of Young’s moduli obtained for the BAPN treated mouse pancreatic ductal adenocarcinoma KPC tumor model (BAPN) using AFM indentation technique with maximal indentation depth by AFM probe equal 2µm. Distributions were fitted with probability density function of the log normal distribution and corresponding mean ± SD values are presented on the right side of each distribution.

**Figure S9.**
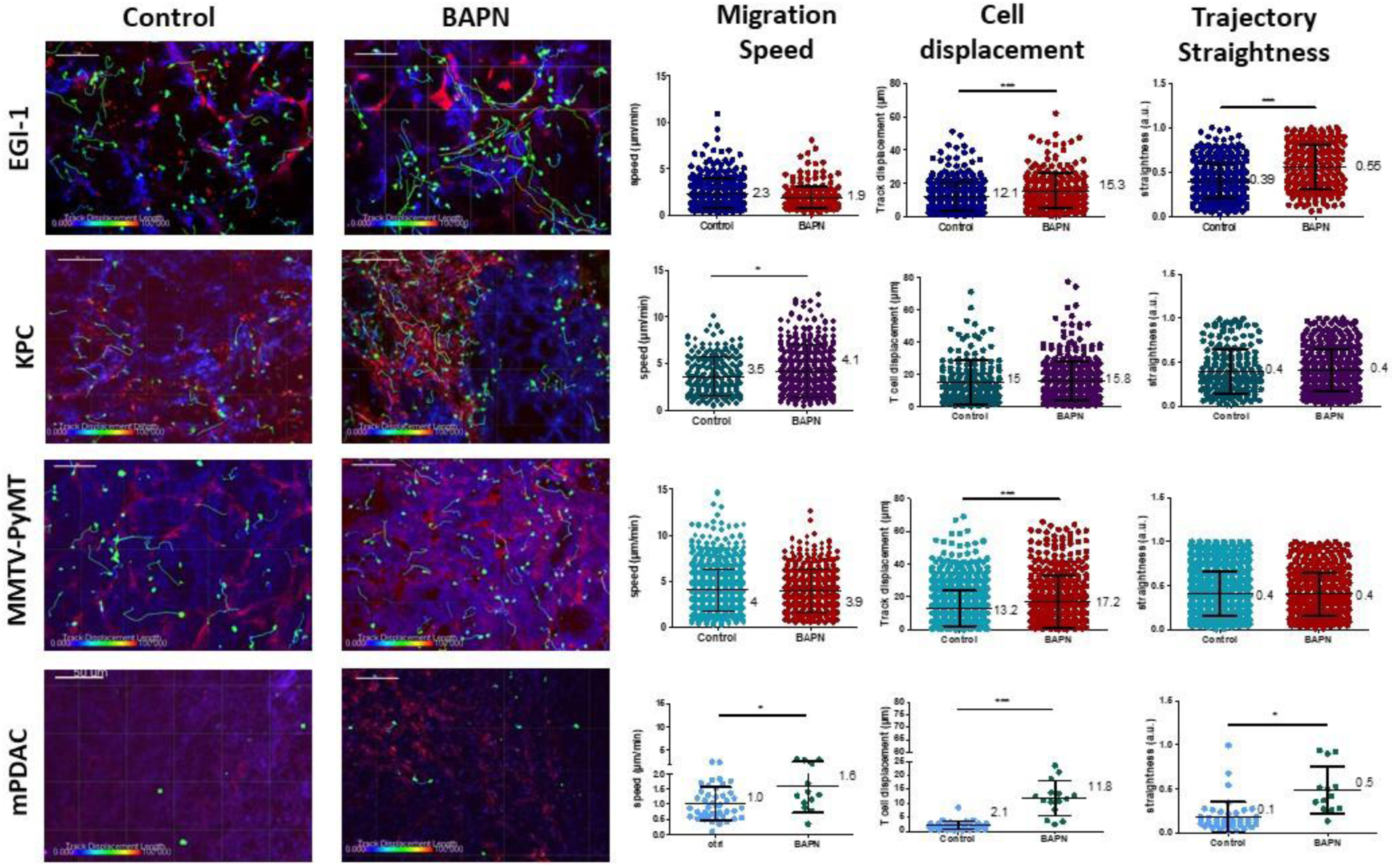
Impact of LOX inhibition on T cell migration in EGI-1, MMTV-PyMT, mPDAC and KPC tumor models. Migration of activated PBT plated onto fresh tumor slices was analyzed in EGI-1 and MMTV-PyMT tumor model, whilst resident tumor-infiltrating T lymphocytes were analyzed in mPDAC and KPC tumor model. Illustrative images of T cell migration tracks in EGI-1, MMTV-PyMT, mPDAC and KPC tumor models. Tumor stroma (fibronectin) in red, tumor cells (EpCAM in EGI-1, MMTV-PyMT and KPC tumor models, CD44 in mPDAC tumor models) in blue and T cells (CD8 in mPDAC and KPC, Calcein in MMTV-PyMT and EGI-1 tumor models) in green. Tracks are color coded to illustrate track displacement. Scale bar = 100 µm. T cell migration speed, T cell displacement and trajectory straightness in all tumor models. *** *P-*value > 0.001, *P-*value > 0.05, Student t test. Results are shown as mean ± SD.

**Figure S10.**
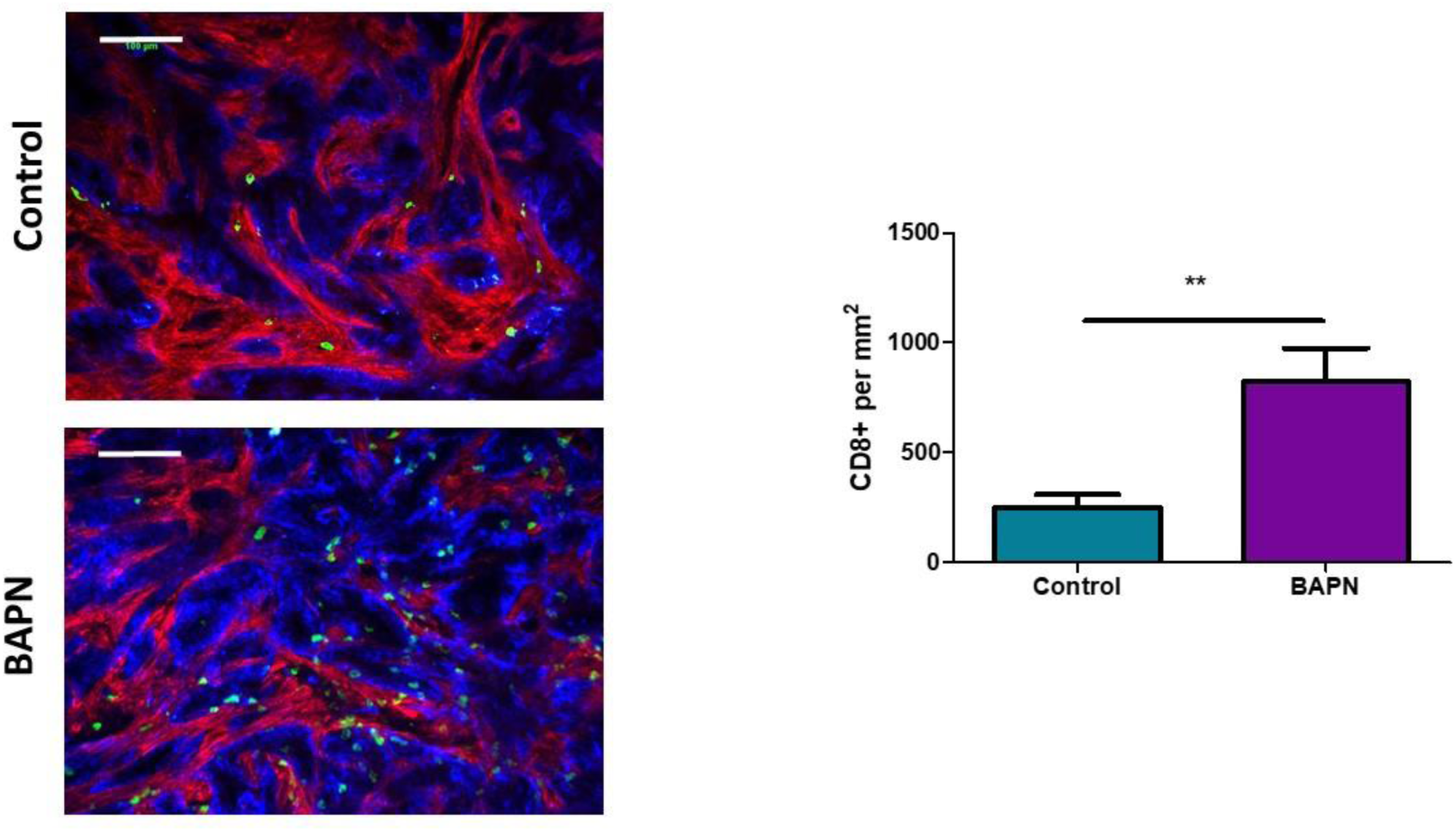
BAPN treatment induces an increased infiltration of effector CD8 T cells in KPC tumors. Representative images of Control and BAPN treated KPC tumors. Tumor cells (EpCAM) in blue, tumor stroma (fibronectin) in red and T cells (CD8) in green. Scale bar = 100 µm. Quantification of CD8+ T cell per mm2 of tissue. (** P < 0.01, Student t test, n = 5/ group).

**Table S1.**
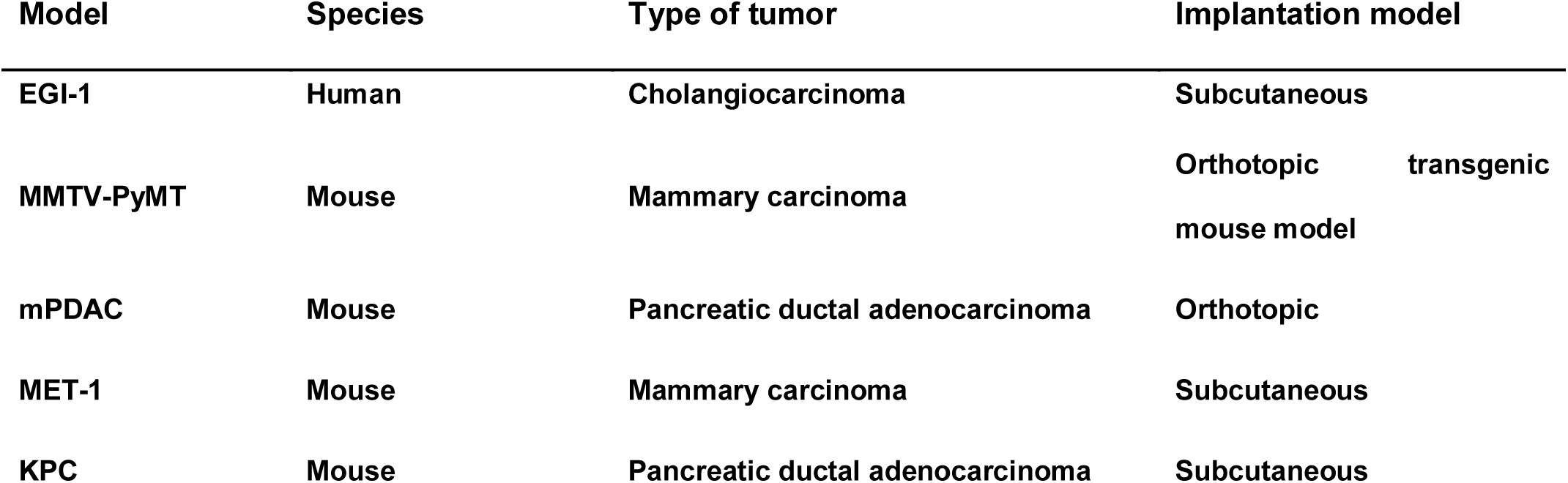
Summary of relevant preclinical models for solid tumors used in this study.

**Table S2.**
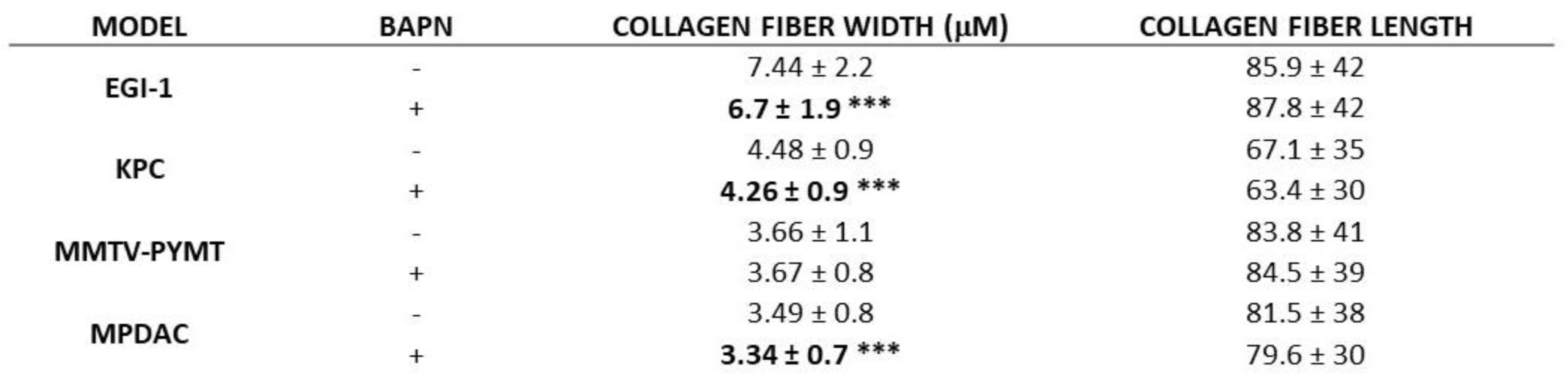
Collagen fiber width and length in EGI-1, MMTV-PyMT, mPDAC and KPC tumor models in control and treated conditions. Results are shown as mean ± SD.

#### Supplementary Movies

**Movie S1. T cell migration in the control EGI-1 tumor model.** The slices stained with calcein for the T cells (green), anti-EpCAM (blue), anti-fibronectin (red) were imaged with a spinning disk confocal microscope. Tracks are color-coded according to T cell displacement length. Frame interval 30s. The animations represent 3D reconstructions of sequential z series. A still image is shown on Table 2.

**Movie S2. T cell migration in the LOX inhibited EGI-1 tumor model.** The slices stained with calcein for the T cells (green), anti-EpCAM (blue), anti-fibronectin (red) were imaged with a spinning disk confocal microscope. Tracks are color-coded according to T cell displacement length. Frame interval 30s. The animations represent 3D reconstructions of sequential z series. A still image is shown on Table 2.

## SUPPLEMENTAL EXPERIMENTAL PROCEDURES

### T cell preparation

Human CD8+ T cells were isolated from cytapheresis rings (obtained from Establissement Français du Sang) using EasySep™ Human CD8+ T Cell Isolation Kit (Stem Cell Technologies), following manufacturers protocol. They were cultured in RPMI supplemented with 3% Human AB serum, T cell TransAct (Miltenyi Biotech), 155 U/mL of human IL-7 (Miltenyi Biotech) and 290 U/mL of human IL-15 (Miltenyi Biotech). 2-3 after activation the cell culture media was changed using the same recipe without TransAct. At day 7 after activation T cells were used for migration experiments.

Mouse CD8+ T Cells were isolated from FVB mouse spleen and lymph nodes using CD8□+ T cell Isolation Kit (Miltenyi Biotech) following manufacturers protocol. Isolated T cells were activated using the Dynabeads™ Mouse T-Activator CD3/CD28 for T-Cell Expansion and Activation (Thermo Fischer) following manufacturers protocol. At day 7 after activation T cells were used for migration experiments.

### Mouse tumor models

#### EGI-1 subcutaneous model

2×10^6^cells suspended in 60 μL of PBS were mixed with 60 μL of Matrigel® growth factor reduced (Corning) and implanted subcutaneously into the flank of 5-week-old female NMRI-nu (nu/nu) mice (Envigo, France).

#### KPC subcutaneous model

3×10^6^cells suspended in 50 μL of PBS were mixed with 50 μL of Matrigel® growth factor reduced (Corning) and implanted subcutaneously into the flank of 6-week-old female C57BL/6J mice (Janvier, France).

#### MET-1 subcutaneous model

10^6^cells suspended in 50 μL of PBS were injected subcutaneously into the flank of a 6-week-old female FVB mice (Janvier France).

#### MMTV-PyMT model

FVB MMTV-PyMT model is maintained at the Cochin Institute specific-pathogen-free-animal facility in accordance with the University Paris Descartes ethical guidelines.

#### mPDAC orthotopic model

10^3^cells suspended in 50 μL of PBS were injected orthotopically in the pancreas of 6-week-old FVB/n mice (Janvier, France). Tumor growth was followed through ultrasound imaging using a VEVO2100 (Visualsonics).

### Shear wave elastography

The mice were anesthetized with 2% isoflurane and their body temperature was maintained at physiological level using a heating plate. B-mode images and SWE images were acquired simultaneously. The B-mode image allowed us manually determine the region of interest (ROI) corresponding to the tumor contours. SWE mode was performed using the penetration mode with a color scale ranging from 0 (transparent) to 40 kPa (red), arbitrarily chosen in the beginning of the study according to the expected stiffness values. The area, the diameter and a set of stiffness values (mean, minimum, maximum and standard deviation) were recorded for the ROI as previously defined. SWE images were also analyzed using an in-house MATLAB code to recover the stiffness map.

### Imaging of CD8 T cell motility

To evaluate T cell migration in EGI-1 model, activated human CD8+ T cells were first labelled with calcein orange red dye (Thermo Fischer). Briefly, T cells were incubated with 125 nM Calcein in HBSS solution for 10 minutes at 37°C. The staining reaction was stopped by adding cold HBSS supplemented 2% SAB. Cells were then pelleted and diluted to an appropriate concentration in phenol red free RPMI media. 2×10^5^cells suspended in 50 μL of phenol red free RPMI media and added on 350µm tumor slices placed on top of 0.4-μm organotypic culture inserts (Millicell; Millipore) in 35-mm petri dishes containing 1.1 mL RPMI 1640 without Phenol Red. The tumor slices were then incubated for 30 min at 37°C and 5% CO_2_. The slices were then washed to remove all cells that had not infiltrated the slice and stained for 15 min at 37 °C with the following antibodies: BV421–anti-human EpCAM (9C4 clone; BioLegend) and anti-human/mouse fibronectin at 10µg/mL. The same protocol was followed to evaluate T cell migration in MMTV-PyMT cells except that activated murine CD8+T cells were used.

To evaluate resident T cell migration in KPC and mPDAC model, tumor slices were then transferred to 0.4-μm organotypic culture inserts (Millicell; Millipore) in 35-mm petri dishes containing 1.1 mL phenol red free RPMI media. Live vibratome sections were stained with BV412 anti-mouse EpCAM (G8.8 clone; BD Biosciences), PerCP-e710 anti-mouse CD8a (53-6.7 clone, eBioscience), PE anti-podoplanin (8.1.1 clone; BioLegend) and anti-human/mouse fibronectin.

T cells were imaged with a DM500B upright microscope equipped with an upright spinning disk confocal microscope (Leica) equipped with a 37 °C thermostatic chamber. For dynamic imaging, tumor slices were secured with a stainless steel slice anchor (Warner Instruments) and perfused at a rate of 0.8 mL/min with a solution of RPMI without Phenol Red, bubbled with 95% O2 and 5% CO2. Ten minutes later, images from a first microscopic field were acquired with a 25× water immersion objective (20×/0.95 N.A.; Olympus). For four-dimensional analysis of cell migration, stacks of 10–12 sections (z step = 5 μm) were acquired every 30 s for 20 min at depths up to 80 μm. Regions were selected for imaging when tumor parenchyma, stroma, and T cells were simultaneously present in the same microscopic field. For most of the tumors included in the study, between two and four microscopic fields were selected for time-lapse experiments.

### Second harmonic generation microscopy

The laser beam was circularly polarized and a Leica Microsystems HCX IRAPO 25×/0.95 W objective was used. Second harmonic generation (SHG - collagen structure) signal was detected in epi-collection through a 405/15nm bandpass filters respectively, by NDD PMT detectors (Leica Microsystems) with a constant voltage supply, at constant laser excitation power, allowing direct comparison of SHG intensity values. All images were then analyzed using CT-FIRE software (*53*) to obtain the width and the length of the collagen fibers. To calculate the curvature ratio, line regions were drawn along the length of the fibers (A) and along the linear distance between the start and the end of the fibers (B), the curvature ratio was calculated CR = A/B. For each image at least 15 fibers were analyzed. Fiber alignment was determined using the directionality plugin in Image J. The fiber alignment was defined by the coefficient of variation (CV) of the angle for all fibers per image. The smaller the CV is, the more aligned the fibers are.

### Histology

After fixation, tumors were dehydrated in graded solutions of ethanol and embedded in paraffin. Five micrometer tissue sections were stained with hematoxylin / eosin / Saffron (HES) and Sirius Red staining. For Sirius red stained slices, linear polarized light or brightfield microscopy was performed using full field microscopy (Statif Axio Observer Z.1, Zeiss) equipped with a linear polarizer and a 20x dry objective (Plan Achromatic (NA = 0.7). Under polarized light thin fibers show a greenish-yellow birefringence whilst thicker and densely packed fibers give an orange red birefringence. The percentage of Sirius red staining defined the amount of thick and densely packed fibers, to do so images were split in the RGB channel and the signal in the red channel was quantified.

### Atomic Force Microscopy and shear rheometry

AFM experiments were made maximally 3 hours after sample thawing and tissues were stored and kept in culture medium during experiment at room temperature. Force vs. indentation curves were collected using a silicon nitride cantilever with a spring constant of 0.6 *N*/*m* and a 4.5 μm diameter bead attached. Each sample was indented in the multiple locations to account for possible heterogeneity in tissue mechanical properties. Briefly, up to nine maps consisting of 64 points corresponding to the scan area of 10×10 µm (for one map) per sample were made. Final Young’s modulus values were derived from the Hertz-Sneddon model applied to force vs. indentation curves (*1*) assuming the spherical shape of the probe and Poisson’s ratio equal 0.5. Distributions of the Young’s modulus values for each control and treated sample, as well as the mean values along with standard deviations were prepared.

For the macroscopic rheological tests HAAKE Rheostress 6000 rheometer (Thermo Fisher Scientific, Waltham, MA, USA), fitted with 8 mm diameter parallel plate system was used. Tissues were firstly cut into disk-shaped samples using 8 mm punch. To avoid tissue slippage during the tests, samples were arranged in sand paper glued inside the Petri-dishes and fixed to the rheometer bottom plate. Rheological experiments were made maximally 3 hours after tissue thawing, and all the samples were kept in humid conditions during the experiment. The rheological evaluation consisted of the oscillating shear deformation with 2% shear amplitude andd frequency of 1 Hz. Final results are presented as the mean values of storage modulus (*G*’) of control and treated tumors ± standard deviation (SD) values.

### Flow cytometry

The anti-mouse antibodies used were the following:

| Antigen | Fluorophore | Clone | Company |
| --- | --- | --- | --- |
| CD11b | PE-Cy7 | M1/70 | Biolegend |
| CD11b | BV421 | M1/70 | BD Biosystems |
| CD11c | BV605 | N418 | Biolegend |
| CD45 | AF700 | 30-F11 | Biolegend |
| Ly-6C | APC-Cy7 | HK1.4 | Biolegend |
| Ly-6G | BV510 | 1A8 | Biolegend |
| CD4 | BV 711 | GK1.5 | BD Biosystems |
| CD8a | PerCP-e710 | 53-6.7 | eBioscience |
| TCRβ | BV605 | H57-597 | BD Biosystems |
| Nkp46 | eFluor660 | 29A1.4 | eBioscience |
| PD-1 | BV421 | 29F.1A12 | Biolegend |
| MHC II | BV785 | M5/114.15.2 | Biolegend |
| CD80 | FITC | 16-10A1 | Biolegend |
| CD64 | PE | X54-5/7.1 | Biolegend |
| CD206 | BV605 | CO68C2 | Biolegend |
| F4/80 | BV650 | A3-1 | Biolegend |

After surface staining, cells were fixed with BD Fixation and Permeabilization Solution for 20 min at 4 °C. In the case of intracellular staining, cells were incubated overnight with the following anti-mouse antibodies:

| Antigen | Fluorophore | Clone | Company |
| --- | --- | --- | --- |
| FoxP3 | FITC | MF-14 | Biolegend |
| GrzmB | PE | QA16A02 | Biolegend |

After washing in PBS, cells were resuspended in PBS 2% FBS and analyzed with a BDFortessa flow cytometer (BD Bioscience). Data were analyzed by FlowJo software.

